# Investigation of CD36 interactome provides insights into multimolecular complexes necessary for anti-angiogenic signalling

**DOI:** 10.1101/2025.07.15.664851

**Authors:** Arashdeep Saini, Erik Gomez-Cardona, Han Huang, Swai Mon Khaing, Katie Klein, Khuloud Jaqaman, Olivier Julien, Nicolas Touret

## Abstract

Signal transduction is a fundamental process that enables cells to adapt to external cues and organize adequate responses including survival, death, growth, and homeostasis. A key mechanism modulating signal transduction relies on the formation of multimolecular complexes optimized for specificity, modularity and signal amplification. The scavenger receptor CD36, which binds diverse ligands in different cellular contexts, illustrates this principle. To uncover the nature of CD36 multimolecular complexes, we employed a proximity biotinylation labeling approach on human endothelial cells, where CD36 binds to thrombospondin-1 (TSP-1) to initiate a signaling cascade promoting programmed cell death. Using biotin capture and mass spectrometry protein identification, we uncovered a list of proteins in the vicinity of CD36. This list of candidates was refined by proximity ligation assays. The relationship between key CD36 interacting molecules, in particular active integrin beta-1 (ITGB1) and CD9, was further decoded by conditional colocalization analysis, providing support for their association within a tri-molecular complex. The implication of selected candidates in the signaling function of CD36 was further evaluated using shRNA knockdown, revealing that active ITGB1 is essential for Fyn activation downstream of CD36, with the tetraspanin playing a connecting role between CD36 and active ITGB1. Our approach to investigating CD36 complexes emphasizes the complexity and fundamental role of protein-protein interactions and coordination in the context of transmembrane signal transduction.

## Introduction

A key mechanism modulating signal transduction is receptor clustering, in which membrane receptors aggregate into complexes often facilitating permissive multimolecular interactions. This spatial arrangement influences signaling efficiency via increased avidity, specificity and signal amplification (Cebecauer et al., 2010; Bethani et al., 2010; Garcia-Parajo et al., 2014; Garcia-Parajo and Mayor, 2024). While this phenomenon has been best characterized for immune receptors such as Fc, T and B cell receptors (Li and Yu, 2021; Treanor et al., 2010; Wilson et al., 2011; Goyette et al., 2019), it is likely a much broader and under-investigated process (due to limitations in identification and analysis of these nanoscale multimolecular complexes) (Chen et al., 2021; Suzuki et al., 2007; Kalappurakkai et al., 2018; O’Shea and Murray, 2008). The nature of these signaling platforms could be modular and cell specific, thus explaining how receptors signaling varies in different contexts. The scavenger receptor CD36 is a typical example of such modularity.

CD36 is a class B scavenger receptor that is expressed primarily at the surface of macrophages, platelets, microvascular endothelial cells, adipocytes, and myocytes (Febbraio et al., 2001; Silverstein and Febbraio, 2009). CD36 has a predominant extracellular loop that contains binding domains for various ligands, two transmembrane domains and two very short intracellular regions lacking scaffolding or signaling motifs. CD36 has been shown to transduce signaling cascades when binding various ligands depending on the cellular context, such as ß-amyloid on the surface of microglia cells (Bamberger et al., 2003; Wilkinson et al., 2006), oxidized LDL on macrophages (Collins et al., 2009; Jaqaman et al., 2011; Chen et al., 2015; Heit et al., 2013; Wong et al., 2016), thrombospondin-1 (TSP-1) on endothelial cells (ECs) (Chen et al., 2000a; Jiménez et al., 2000a; Githaka et al., 2016), malaria-infected erythrocytes to ECs and other blood circulating cells (Ockenhouse et al., 1989; McGilvray et al., 2000; Davis et al., 2012). CD36 also acts as a fatty acid transporter (Harmon and Abumrad, 1993; Pepino et al., 2014; Chen et al., 2022). One commonality between these ligands is their multivalent characteristic, which emphasizes the requirement for receptor clustering for their activity. Previous studies have shown that CD36 resides in nanodomains, areas within the plasma membrane enriched in proteins and lipids, regulated by the actin cortex, which support signaling and responses to cellular stimuli (Jaqaman et al., 2011; Githaka et al., 2016; Dasgupta et al., 2023). Upon stimulation with multivalent ligands, such as TSP-1 or anti-CD36 IgM (Dawson et al., 1997; Jiménez et al., 2000a; Githaka et al., 2016), the enhancement of CD36 nanoclusters leads to activation of the Src-family kinase Fyn, culminating in EC apoptosis (Dawson et al., 1997; Jiménez et al., 2000a). This anti-angiogenic pathway has been targeted for therapeutic intervention in the treatment of certain cancers (Wu and Finley, 2017; Lawler, 2022; Wen et al., 2024).

Given the lack of intrinsic signaling capacity in CD36 and its reliance on clustering for signal transduction, we hypothesized that CD36 functions within multimolecular complexes, the composition of which is regulated by protein-protein interactions and the membrane ultrastructure. To enhance our understanding of CD36 signaling complexes and their nanoscale organization, we sought to identify proteins implicated in this anti-angiogenic response in endothelial cells by adapting a proximity biotinylation approach developed by Bar and colleagues (Bar et al., 2018).

To discover proteins residing within CD36 nanoclusters, we engineered human telomerase-immortalized microvascular endothelial cells (TIME) to express the receptor in an inducible manner and employed biotinylation by antibody recognition (BAR) proximity labeling followed by tandem mass spectrometry protein identification (Bar et al., 2018). Confirmation of candidate protein proximity via in situ-proximity ligation assay (Söderberg et al., 2006) narrowed our investigation to study the role integrin beta1 (ITGB1) and the tetraspanin CD9. To determine if these candidate proteins were necessary for CD36 anti-angiogenic signaling, we performed CD36 stimulation experiments within TIME CD36 cells where CD9 or ITGB1 where silenced using shRNA. Conditional colocalization analysis further elucidated the relationship between CD36, CD9, and ITGB1, and the alterations in their interactions during CD36-Fyn signaling (Vega-Lugo et al., 2022). Through investigation of the CD36 interactome, we have revealed a CD36, CD9, and ITGB1 complex that is implicated in anti-angiogenic signaling. This provides new insights into membrane protein interactions and organization that could promote signal transduction in an indirect manner, e.g. when the receptor does not contain intrinsic signalling domains or motifs.

## Results

### Identification of CD36 interactors via biotinylation by antibody recognition (BAR)

Our previous work has underlined how the association between CD36, Fyn, membrane lipid domains and actin cortex in ECs are required for signal transduction (Githaka et al., 2016; Dasgupta et al., 2023). Given its short cytosolic domains, and the requirement for clustering, we hypothesized that CD36 must function within multimolecular complexes to transduce signals. To identify proteins residing in proximity of CD36 at the surface of endothelial cells, we employed biotinylation by antibody recognition (Bar et al., 2018) targeting CD36 at the surface of TIME cells (hTERT Immortalized Microvascular Endothelial cells, ATCC CRL-4025) engineered to express mEmerald-CD36 (mEm-CD36) under the control of an inducible *tet*-on promoter.

The method relies on the production by a peroxidase (horseradish peroxidase, HRP) of reactive and short-lived biotin radicals that react to exposed tyrosine residues, usually within a ∼250 nm radius (Bar et al., 2018; Oakley et al., 2022; Zafra and Piniella, 2022) (Fig. 1A). We first validated the efficacy and specificity of the CD36 BAR methodology through detection of biotinylation events at the surface of induced TIME mEm-CD36 cells (0.5 µg/mL of doxycycline for 18h) versus non-induced cells (no CD36 expression) (Fig. 1B). Cells were fixed and labeled using mouse anti-CD36 and goat anti-mouse Fab coupled to HRP antibodies. A Fab fragment antibody was used instead of regular IgG to reduce the size of the antibody complex and limit the distance of the peroxidase from CD36. The biotinylation reaction (see materials and methods) was performed for 10 min at 37°C on cells previously fixed on ice. Protein biotinylation events were detected using streptavidin coupled to Alexa Fluor 647 (Streptavidin-AF647) and imaged on a widefield microscope. mEm-CD36 expression (Fig. 1B) was confirmed in cells induced with doxycycline, whereas uninduced cells had no CD36 signal. For BAR negative control treatment on non-induced cells, we observed low-intensity punctate labeling with streptavidin-AF647, indicative of a low level of non-specific biotinylation (Fig. 1B). In contrast, streptavidin-AF647 signal was robust (same imaging and image contrast parameters between conditions) on TIME mEm-CD36 expressing cells compared to control. The overlap between the streptavidin-AF647 and mEm-CD36 signals demonstrated the specificity of the labeling (Fig 1B). Most of the streptavidin-AF647 signal overlapped with only the cell surface fraction of mEm-CD36. Therefore, we proceeded to capture biotinylated proteins on whole cells lysates extracted from similar experimental conditions.

**Figure 1:**
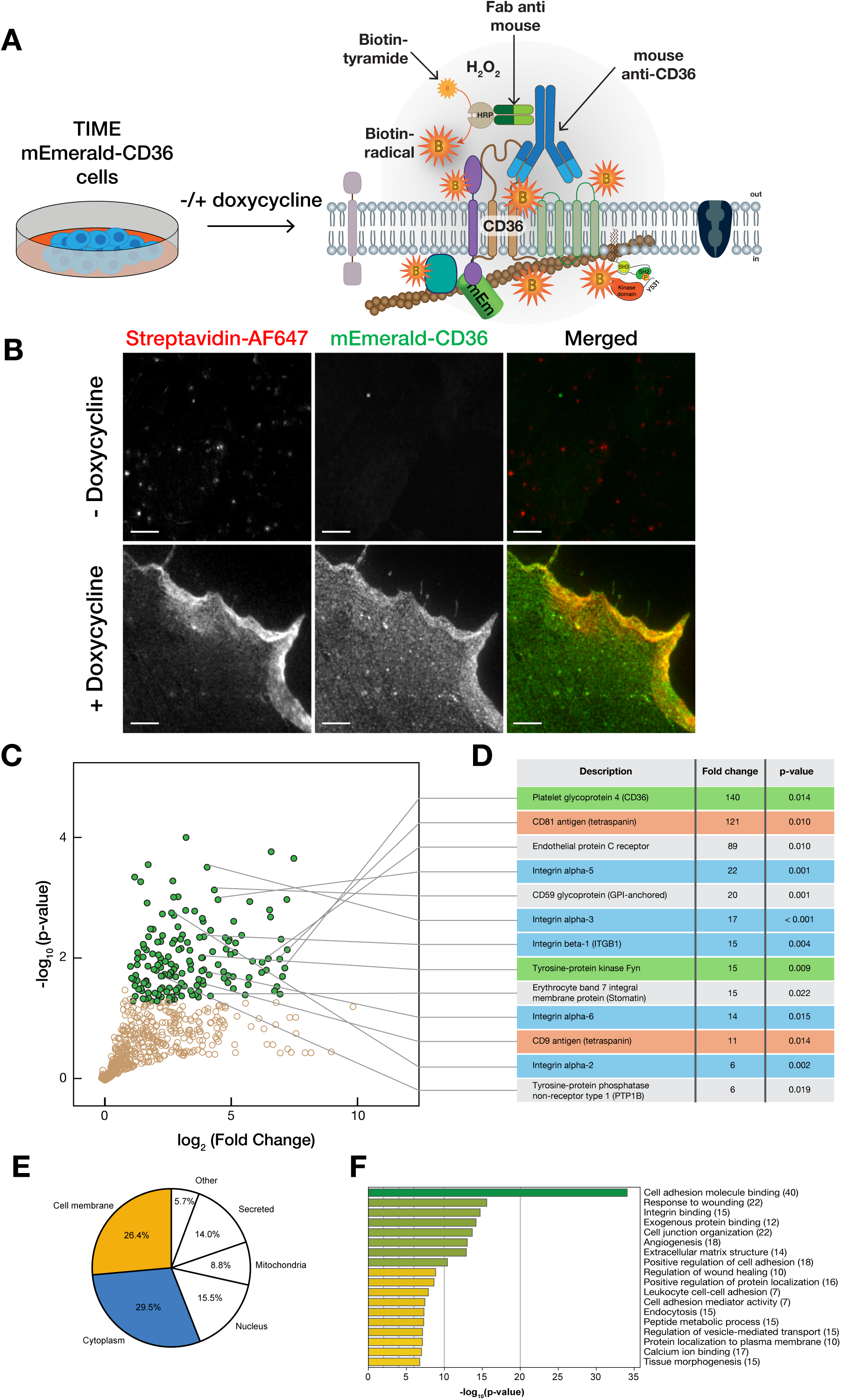
Workflow of CD36 BAR protein methodology and protein classification. (**A**) Schematic of CD36 BAR workflow consisting of inducing or not mEm-CD36 expression on TIME mEm-CD36 ECs and followed by BAR experiment. CD36 is labeled using IgG mouse anti-CD36 (blue antibody) followed by incubation with goat anti-mouse Fab conjugated to HRP (green) antibody. Reactive biotin radicals are generated by HRP from biotin-XX-tyramide and hydrogen peroxide. Proteins within a ∼250 nm radius (symbolized by gray circle) can be covalently labeled with biotin on exposed tyrosine residues. (**B**) Representative images of CD36 and protein biotinylation events for CD36 BAR method. TIME mEm-CD36 cells were either uninduced (BAR control: - doxycycline) or induced with doxycycline (CD36 BAR: + doxycycline). Biotinylated proteins were detected on fixed cells after CD36 BAR treatment using streptavidin-AF647. Scale bar = 5 µm. (**C**) Protein identification using mass spectrometry. Significantly enriched proteins in the CD36 BAR dataset are colored in green. Enrichment threshold was set for ratios (CD36 BAR/BAR control) > 2 (log_2_ (2) = 1), and p-values < 0.05 (-log_10_(0.05) = 1.3). (**D**) A set of candidate proteins were labeled to visualize their position on the volcano plot. Proteins highlighted in red are members of the tetraspanin superfamily, blue are a part of the integrins family, and in green are CD36 and Fyn, expected to be enriched in our CD36 BAR dataset. (**E**) Pie chart displaying the distribution of CD36 BAR enriched proteins based on their subcellular localization. (**F**) Metascape gene analysis indicates the number and the enrichment of proteins within various cellular functions.

To explore both the efficacy of the biotinylation reaction and the efficiency of the biotin capture, we performed detection of biotinylated proteins within whole cell lysates in control cells and cells expressing mEm-CD36 using streptavidin blots (Suppl. Fig.1). After the biotinylation reaction and lysis, ∼10% of the sample was kept as whole cell lysate (WCL), while the remaining 90% was used for biotinylated protein capture using magnetic streptavidin beads in an automated Duo Prime Kingfisher instrument (ThermoFisher Scientifc, Waltman, MA, USA). Flow-through (FT) and WCL from 8 experiments were analysed by SDS-PAGE, transferred to nitrocellulose membrane and probed with fluorescent streptavidin (streptavidin-IRDye 800RD) (Suppl. Fig.1). Biotinylated proteins were detected in CD36 BAR lysate, whereas little to no biotinylated proteins were visualized in the CD36 BAR control lysate confirming the specificity of the CD36 BAR method. The strong signal in the WCL lanes confirmed efficient biotinylation, and the lack of signal on the flow-through lanes indicated efficient capture by the streptavidin beads (Suppl. Fig. 1).

The captured proteins were ‘on-beads’ trypsin digested and processed for liquid chromatograph followed by tandem mass spectrometry (LC-MS/MS). Approximately a thousand proteins were identified in each experiment, regardless of CD36 expression (Table 1). To compare quantitatively each experimental condition, we designed a normalization strategy (see Materials and Methods and Suppl. Fig. 2) that used the mean intensity of a set of common proteins found in each condition as the normalizing factor. The normalized protein intensities were averaged, and statistical comparisons were performed between CD36 BAR and BAR control conditions. An enrichment ratio was calculated for each protein as the average intensity for CD36 BAR divided by its average intensity for BAR control. The results confirmed the specificity of the CD36 BAR technique as the abundance of CD36 itself is ∼140-fold greater in CD36 BAR than in BAR control (Fig. 1 C and D). Known downstream effector of TSP-1 and CD36, Fyn, was found enriched ∼15 times more in the CD36 BAR samples compared to BAR control (Fig. 1 C and D). Proteins with enrichment ratios greater than 2, and whose average protein intensities in CD36 BAR were statistically different from control, resulted in a list of 153 proteins (Fig. 1C and Suppl. Table 1). Of these proteins, ∼25% were membrane associated (54 proteins, Fig. 1 E) and ∼25% corresponded to cytosolic proteins (Fig. 1E). Although biotin-phenol is impermeable to the cell membrane, intracellular proteins were identified in our CD36 BAR MS/MS dataset. The paraformaldehyde fixation potentially resulted in permeabilization of the cell membrane, allowing biotin radicals to cross the membrane and covalently react with intracellular proteins (Cheng et al., 2019).

**Table 1:**
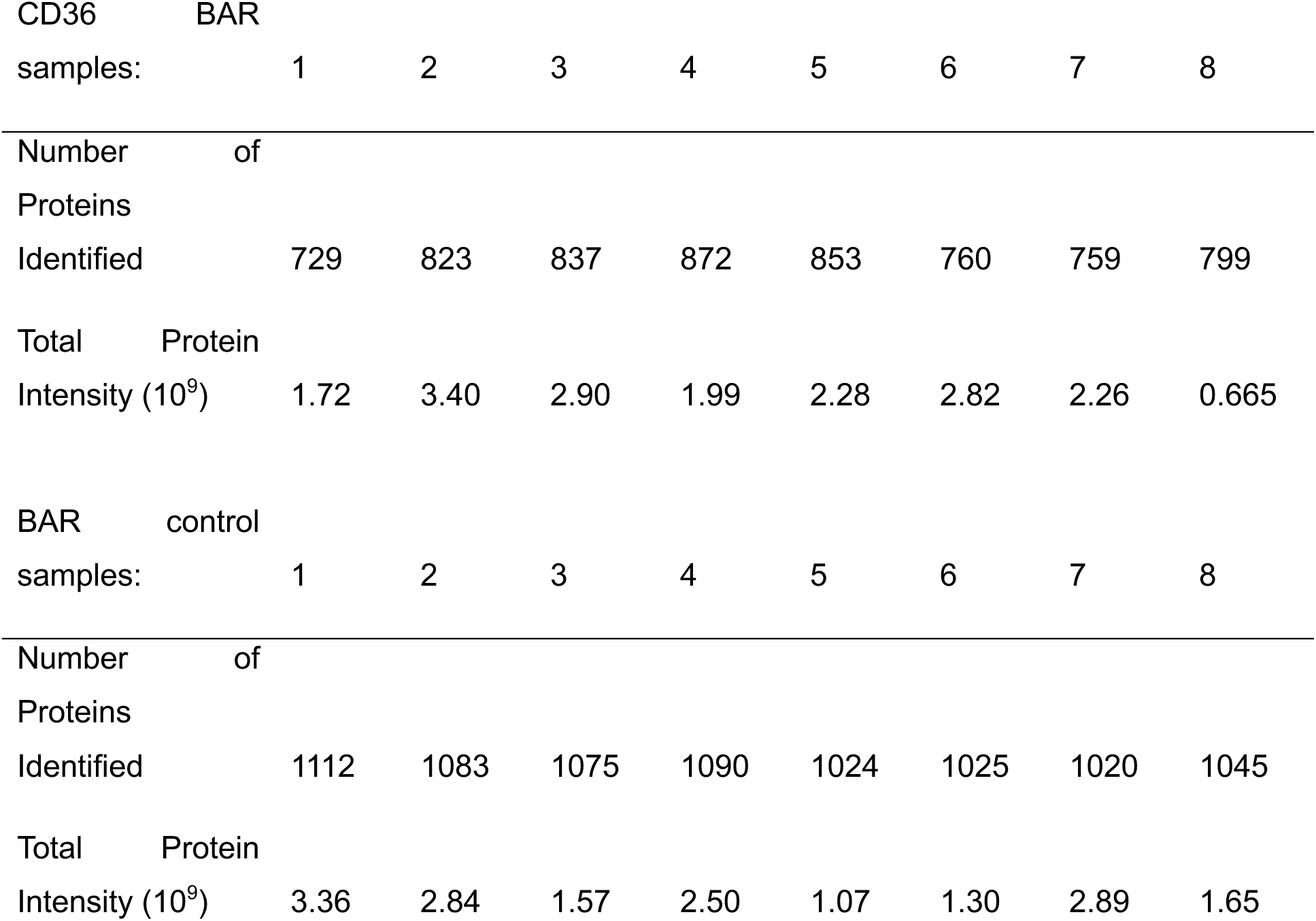
Summary of CD36 BAR experiment results.

To identify possible classes of proteins enriched in the vicinity of CD36, we performed gene annotations analysis on this list using metascape (Zhou et al., 2019). The pathway analysis revealed that amongst the most significantly enriched pathways was angiogenesis, which was expected given the role of CD36 in promoting the TSP-1 anti-angiogenic signaling (Fig. 1F). Interestingly, we also found the enrichment of pathways involving integrin functions such as cell adhesion, integrin binding, and cell and junction organization. In alignment with metascape data, there are 6 proteins of the integrin family and 2 tetraspanins enriched within our CD36 BAR dataset. These data suggest that integrins are near CD36. Our next steps were to confirm the proximity of candidate proteins listed in Fig. 1D to CD36 via in-situ proximity ligation assay (PLA).

### Confirmation of candidate protein proximity via PLA

To confirm the proximity of candidate proteins identified by CD36 BAR (Fig. 1D) we employed an in-situ PLA methodology which detect protein interactions within a ∼40nm distance (Söderberg et al., 2006). We chose 13 proteins to further investigate their proximity to CD36 via PLA. The rationale for selecting these candidates was that these proteins were: 1) shown to be interactors of CD36 in previous studies (CD9 (Miao et al., 2001; Huang et al., 2011; Heit et al., 2013; Huang et al., 2023); CD81 (Heit et al., 2013); CD151 (Kazerounian et al., 2011); ITGB1 (Thorne et al., 2000; Bamberger et al., 2003; Davis et al., 2013; Heit et al., 2013) and potential integrin alpha subunits (ITGA2, ITGA3, ITGA5; stomatin (STOM) (Wu et al., 2022)); 2) known to associate with lipid raft domains (CD59 (Murray and Robbins, 1998; Cross, 2004; Omidvar et al., 2006; Koyama-Honda et al., 2020); STOM (Salzer and Prohaska, 2001; Mairhofer et al., 2002; Umlauf et al., 2006)); and 3) known to have signaling capacity towards Src-family kinases (ITGB1 (Wary et al., 1998; Chen et al., 2000b; Liang et al., 2004; Quintela-López et al., 2019); CD59 (Murray and Robbins, 1998; Suzuki et al., 2007; Koyama-Honda et al., 2020); Tyrosine-protein phosphatase non-receptor type 1 (PTP1B) (Bjorge et al., 2000; Liang et al., 2005; Arias-Romero et al., 2009).

PLA reactions were carried out using one antibody to detect mEm-CD36 in induced TIME mEm-CD36 cells and a second against one of the candidates (see Table 3 and 4). To quantify PLA images, we generated an analysis pipeline in CellProfiler (Carpenter et al., 2006; Stirling et al., 2021) to determine the density of PLA dots per cell (Fig. 2A and Materials and Methods). The density measurement allows normalization to different cell area, and density of dots per cells was compared between all candidates (Fig. 2B). As a positive control, we performed PLA utilizing 2 different primary antibodies directed to 2 different epitopes on mEm-CD36, a mouse anti-CD36 and a rabbit anti-GFP. This produced the highest density of PLA dots and was ∼18 times more than the negative control performed using mouse anti-mitofilin (mitochondrial inner membrane protein) and rabbit anti-GFP (Fig. 2B).

**Figure 2:**
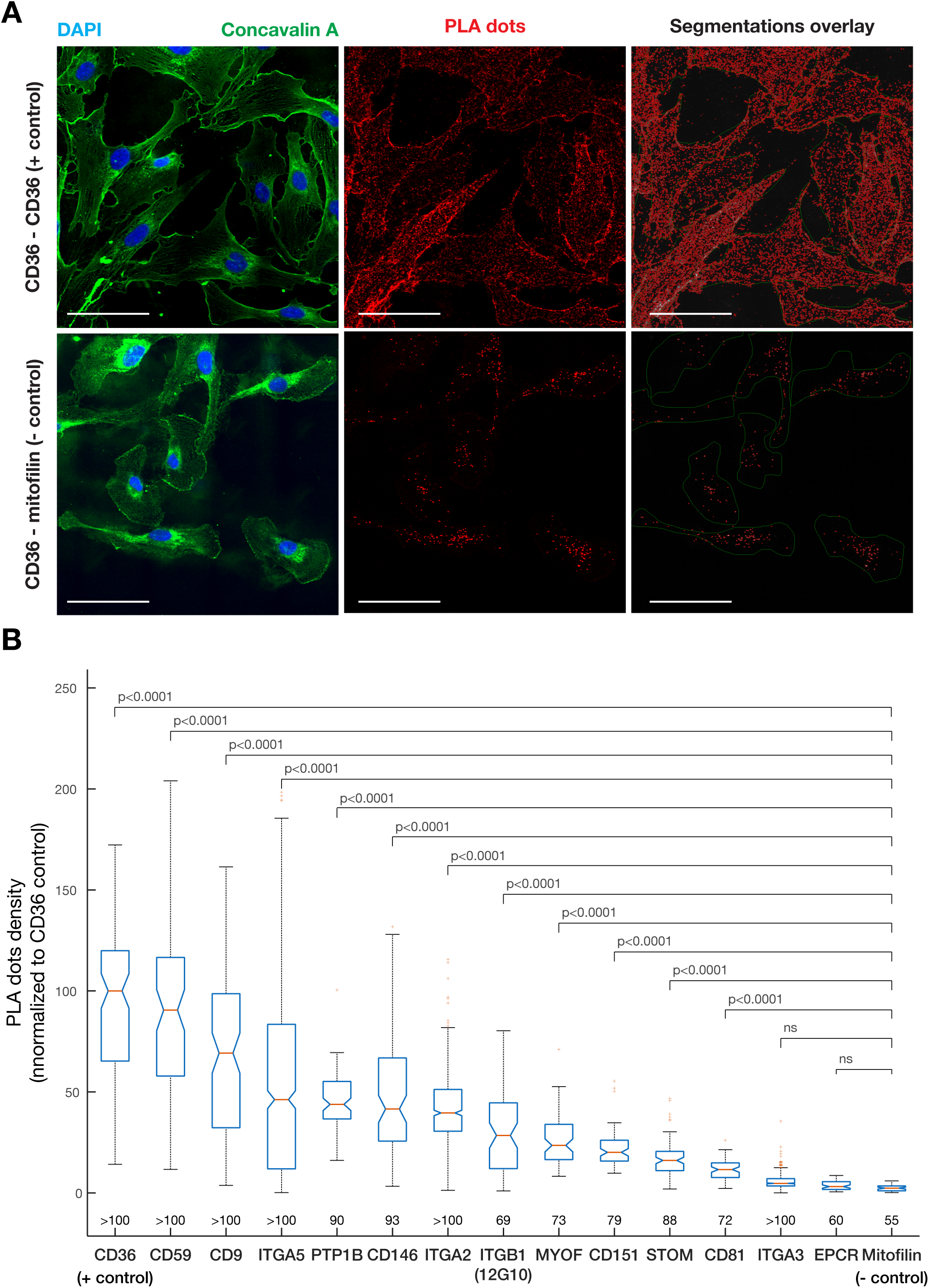
Candidate proteins association with CD36 accessed via PLA. (**A**) TIME mEmerald-CD36 cells were processed for PLA between CD36 and each one of the candidate proteins. PLA dots quantification was performed in Cell Profiler (Carpenter et al., 2006; Stirling et al., 2021) and based on the numeration of the PLA dots within the cells’ area, segmented using the concavalinA-AF488 labeling. Representative images of PLA dots in positive (mEm-CD36 labeled with mouse anti-CD36 and rabbit anti-GFP) and negative (mEm-CD36 and mitofilin labeled with rabbit anti-GFP and mouse anti-mifofilin) control conditions are shown together with the resulting segmented images (right panels). Scale bar: 100µm. (**B**) Quantification of the density of PLA dots between CD36 and candidate proteins. Boxplots display the PLA dot density per cell normalized to the positive control (mouse anti-CD36 and rabbit anti-GFP). The number of cells analyzed for each candidate protein is denoted below each boxplot. Non-parametric Kruskal-Wallis test was employed to determine if there were statistically significant differences between groups. Dunn’s post-hoc test was used to determine pairwise significance values and are shown above the corresponding boxplots. Non-significant (ns) values are for α value > 0.05.

Using this quantification, we were able to confirm the proximity to CD36 of 10 candidates: CD59, CD9, ITGA5, PTP1B, CD146, ITGA2, active ITGB1, myoferlin (MYOF), STOM and CD81 (Fig. 2B). Because of the role of active ITGB1 in the regulation of the actin cortex (Jiménez et al., 2000b; Vicente-Manzanares et al., 2009; Calderwood et al., 2013; De Franceschi et al., 2015) and given that CD36 nanodomain organization and signaling are known to require cortical F-actin (Githaka et al., 2016; Dasgupta et al., 2023), we focused our investigation on the active conformer of ITGB1 using the antibody clone 12G10. PLA analysis revealed that CD36 and active ITGB1 are in proximity to each other in the plasma membrane. Although EPCR and ITGA3 were significantly enriched within CD36 BAR experiments, these candidate proteins were not significantly different than mitofilin. These proteins were possibly identified within the BAR ∼250nm radius, however, they did not have intimate interactions with CD36 as measured by PLA (∼40nm). Conversely, we found that the tetraspanin CD151, known to interact with CD36 (Kazerounian et al., 2011) produced a PLA signal that was significantly higher than that of the negative control, but it was not one of the selected candidate protein from BAR analysis. In the analysis of the CD36 BAR data, CD151 didn’t reach the significant threshold that we applied for selected candidate proteins. This illustrates the need for alternative methodologies when assessing protein-protein interaction data and grant further exploration of our CD36 BAR results.

### Colocalization between CD36 and active ITGB1 and CD9

To further elucidate the role of candidate proteins in TSP-1 - CD36 signaling, we initially chose to focus on CD9 and active ITGB1 with the following rationale. CD9, a protein of the tetraspanin superfamily, has been shown to interact with CD36 via co-immunoprecipitation in numerous cell lines – human dermal microvascular endothelial cells (HDMEC) (Kazerounian et al., 2011), platelets (Miao et al., 2001), and macrophages (Huang et al., 2011; Heit et al., 2013; Huang et al., 2023). While the role of CD36’s interaction with CD9 is unknown in HDMEC and platelets, this interaction is essential for oxLDL uptake and signaling within macrophages (Huang et al., 2011; Heit et al., 2013; Huang et al., 2023). ITGB1 interaction with CD36 has also been previously described in HDMEC (Kazerounian et al., 2011; Davis et al., 2013) and macrophages (Heit et al., 2013), as well as microglia (Bamberger et al., 2003) and melanoma cells (Thorne et al., 2000). Within microglial cells, CD36 interaction with integrin α6β1 forms a complex that binds β-amyloids fibers and induces cytokines and chemokines production (Koenigsknecht and Landreth, 2004; Bamberger et al., 2003). ITGB1 interacts with CD36 in macrophages, and this interaction is dependent on CD9 (Kazerounian et al., 2011). These three proteins are also involved in a complex needed for oxLDL uptake (Heit et al., 2013). Although the ITGB1 interaction with CD36 has been shown to strengthen *plasmodium falciparum*-infected erythrocytes binding to HDMEC cells, the precise role of this interaction in microvascular endothelial cells and anti-angiogenic signaling remains unclear (Davis et al., 2013). Additional evidence that CD36 organization and signaling relies heavily on the actin cortex (Jaqaman et al., 2011; Githaka et al., 2016) narrowed our focus to active form of ITGB1. Therefore, while the literature and our proximity labeling data strongly suggest interaction between ITGB1, CD9 and CD36 in the membrane of microvascular cells, our objective is to identify the role each of these molecules play during CD36 mediated activation of Fyn. Accordingly, we sought a methodology enabling to categorize the various possible complexes that co-exist in the plasma membrane between CD36, CD9, and active ITGB1, and to identify those that potentiate signal transduction.

To further explore the relationship between CD36, and CD9 or ITGB1, we first sought to determine the colocalization extent of these molecules in a quantitative manner, using object-based two-color colocalization analysis (Helmuth et al., 2010; Vega-Lugo et al., 2022). Low level fluorescent antibody labelling enables the algorithm to localize puncta of the labeled molecules at subpixel resolution and determine their colocalization, defined as occurrence of two labeled molecule types less than 2 pixels (= 180 nm) away (Fig. 3A). TIME Halotag-CD36 (HT-CD36) cells were labeled with 0.25 nM of HaloTag ligand HT-JFX-549 for 10 min at 37°C before fixation on ice and labeling with antibodies already coupled to fluorescent dyes: anti-CD9 AF488 and anti-active ITGB1 (clone 12G10) AF647. Cells were imaged by total internal fluorescence microscopy (TIRFm) as illustrated on Fig. 3A. Images were processed for colocalization analysis, which consisted of segmentation of the cell area, particle (i.e. labeled molecule) localization (see grayscale images with indicated position of detected molecules, Fig. 3A) and analysis of their colocalization relationships as shown on the right panel of Fig. 3A (Vega-Lugo et al., 2022). To estimate the expected range of colocalization between these molecules, we conducted a positive control in TIME HT-CD36 cells by labeling HT-CD36 with 2 different labels. One color was with mouse anti-CD36 (1:5000) and anti-mouse AF488, and the second was HT-JFX 549. For negative control, HT-CD36 labeled with HT-JFX549 was compared to immunostained transferrin receptor (TfR) (refer to Table 5 for antibody concentration). To measure CD36’s overall colocalization with itself or ITGB1, CD9 or TfR, we utilized the *p*(*TwR*) measurement – which determines fraction of CD36 objects (Target (T), HT-JFX549 labeled) colocalizing with antibody labeled HT-CD36, active ITGB1, CD9 or TfR (as Reference (R)) (Fig. 3B). To test the significance of this overall *p(TwR)* measurement, it was compared to the colocalization extent obtained when the positions of target molecules (T) were replaced with point in a grid (nullTR, Fig. 3C) (Helmuth et al., 2010). This value provides the coincidental colocalization reflecting the density and distribution of the reference (R). If not significantly different from the experimental data, then the overall colocalization extent is deemed insignificant. Since CD36 colocalizations with itself, ITGB1, CD9 and TfR are significantly higher than their nullTR counterparts (Fig. 3B), it indicated meaningful interactions.

**Figure 3:**
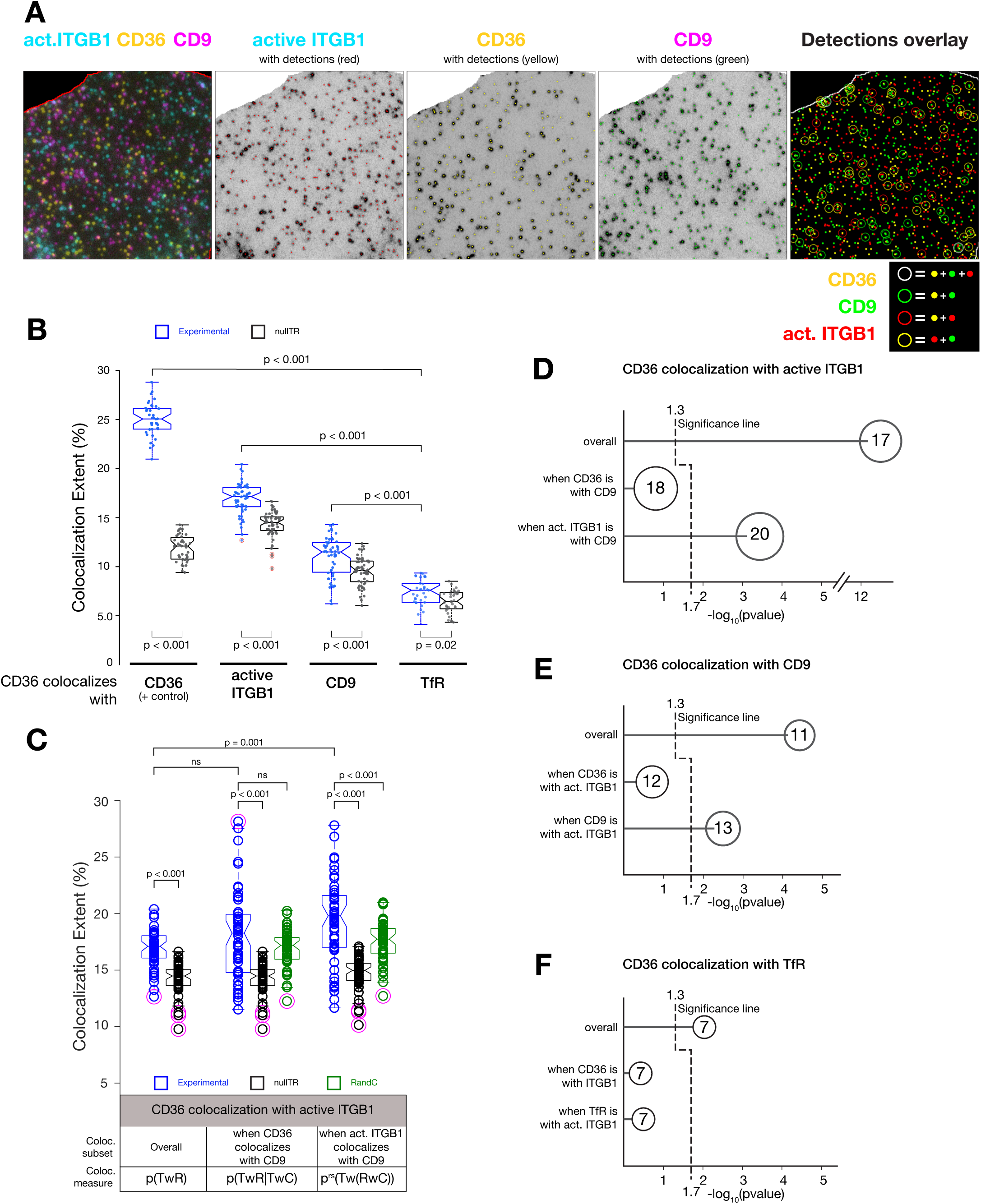
Conditional colocalization analysis between CD36, active ITGB1 and CD9. (**A**) Representative three-color TIRFm images of CD36 (yellow), active ITGB1 (Cyan) and CD9 (Magenta). The coordinates of molecules present in each channel are determined using a point source detection algorithm based on u-track (Jaqaman et al., 2008) (see details in material and methods). After object detection, analysis was performed with a colocalization radius of 2 pixels (180 nm). Triple colocalization events, where CD36, active ITGB1 and CD9 objects were all found within a 2 pixels radius of one another are denoted in white. Two-color colocalization events between CD36-active ITGB1, CD36-CD9, and active ITGB1-CD9 are denoted by red, green and yellow circles respectively. Further statistical analysis from the conditional analysis package (Vega-Lugo et al., 2022) extracts the colocalization extent between 2 molecules and the effect of a third one on their association. (**B**) Two-molecular colocalization measures between CD36 and active ITGB1, CD9 and TfR (as a negative control). Extent of the colocalization of HT-CD36 (labeled with HT-JFX-549) with CD36 (using mouse anti-CD36), or with active ITGB1, CD9 and TfR are shown as boxplots with the experimental data in blue and the nullTR (where (T) positions are replaced with point in a grid) in black. Statistical analysis between experimental data and nullTR was calculated using non-parametric Kruskal-Wallis test with significant difference indicated by p < 0.05. Comparison between groups was measured using non-parametric Kruskal-Wallis test followed by a Dunn’s post-hoc test to determine pairwise significance values, indicated for the relevant comparisons. Results are from 3 replicates and a minimum of 35 cells imaged. (**C**) Conditional colocalization analysis measuring the effect of CD9 on the colocalization of CD36 with ITGB1. The analysis provides a set of conditional colocalization extent measures to establish in a robust statistical fashion the relationship between 3 molecules. Here, the effect of CD36 colocalization with CD9 on CD36’s colocalization with ITGB1 and the effect of ITGB1’s colocalization with CD9 on CD36’s colocalization with ITGB1 are presented. Triple colocalization values were compared to the coincidental colocalization measures (nullTR in black and RandC in green) and overall colocalization measure p(CD36wITGB1) to determine CD9’s colocalization effect on CD36-active ITGB1 colocalization. (**D**) Summary of conditional colocalization measurements from panel C indicating the effect of CD9 on the colocalization between CD36 and ITGB1. Median colocalization extent for each measurement is indicated within each circle. P-values determination for overall colocalizations was done using non-parametric Kruskal-Wallis test between experimental data and nullTR with -log_10_(p-value) above 1.3 indicative of p<0.05. For three-molecular conditional colocalization measurements, the least significant p-value among three comparisons (overall colocalization, nullTR and RandC) was expressed as a -log_10_(p-value). Thresholds for three molecular colocalization significance were calculated with Dunn-Sidak correction reducing the significance threshold to 1.7. (**E**) Similar summary of conditional colocalization determining the effect of active ITGB1 on the colocalization of CD36 with CD9. (**F**)Negative control conditional colocalization values measuring the effect of active ITGB1 on the colocalization of CD36 with TfR. Results are from at least 3 experiments and > 40 cells imaged per condition.

The colocalization analysis of CD36 with itself or with TfR established the upper and lower limits of the colocalization extent values expected with this approach (Fig. 3B). The upper limit reached ∼25% when CD36 was labeled with an antibody and the HT-JFX-549. Even though CD36 molecules were detected with 2 labels, only a fraction of each label is overlapped with the other label. This reflects the specific labeling conditions utilized for this analysis, which relies on identifying individual puncta and therefore requires relatively sparse labeling of a fraction of the total molecules. An overall colocalization extent of ∼25% means that among the molecules labeled with HT-JFX-549, about a quarter also got labelled with the antibodies. For the lower limit, ∼7% of CD36 molecules were found in the vicinity of TfR, which was very close to the coincidental colocalization fraction. CD36 exhibited greater colocalization with active ITGB1 (∼17%) and CD9 (∼11%) than with TfR (Fig. 3B, C), consistent with the interactions identified by proximity biotinylation and PLA analysis.

What really reflects the level of colocalization of two labeled molecules is how significant is the difference between the colocalization extent of the experimental data (*pTwR*) compared to the coincidental nullTR measurement. This comparison revealed that CD36-active ITGB1 and CD36-CD9 colocalizations (pTwR) were ∼1.2 fold higher than nullTR, both passing the significance test with p-values < 0.001. The greater colocalization with active ITGB1 compared to CD9 contrasts with the results of PLA experiments between the same molecules, where the PLA signal was stronger for CD9 than for ITGB1 (Fig. 2B). This disparity is most likely due to variations in antibody concentrations needed for conditional colocalization and PLA experiments. For conditional colocalization, sparse, punctate labelling is required for object detection, whereas PLA requires strong antibody labelling to enhance the connection between oligomer hybridization events. Because of the high proportion of proteins labeled in PLA, proteins which are in proximity to CD36, and with greater expression levels produce more PLA signals. Therefore, since CD9 has ∼2-fold greater expression in TIME cells than ITGB1 (data from The Human Protein Atlas (https://www.proteinatlas.org/) for TIME cells), the PLA produces a more robust signal. The sparse labelling for conditional colocalization nullifies the expression effect, providing a relatively normalized approach to investigate protein interaction. Nonetheless, both PLA and colocalization analysis results confirmed the close relationship that exists between CD36, active ITGB1 and CD9.

### Relationship between CD36, active ITGB1 and CD9

To determine whether CD36, ITGB1 and CD9 form a ternary complex, we next investigated three-molecular colocalization measurements using the conditional colocalization analysis developed by Vega-Lugo et al. (Vega-Lugo et al., 2022). In addition to the two-molecule analysis described above, the conditional colocalization determines whether their colocalization is positively or negatively influenced by the presence of a third molecule type within the same radius (Fig. 3A). Therefore, it is possible to determine whether CD9 and ITGB1 affect one another’s colocalization with CD36. To define the effect of CD9 on the colocalization of CD36 and ITGB1, CD36 was denoted as the target (T), active ITGB1 as the reference (R), and CD9 as the condition (C). The effect of active ITGB1 was conversely determined by setting CD36 as the target (T), CD9 as the reference (R), and active ITGB1 as the condition (C) (Fig. 3C).

For a condition protein (C) to have a significant effect on the colocalization between two proteins (Target with Reference annotated TwR), the three-molecular colocalization measure focused on the subset of target objects colocalized with condition objects (p(TwR|TwC)) and/or the three-molecular colocalization measure focused on the subset of reference objects colocalized with condition objects (p^rs^(Tw(RwC))) must be significantly greater than the two-molecular colocalization measure *p(TwR)* and their respective coincidental colocalization measurements, nullTR (target positions as points on a grid, as above) and RandC (randomized condition positions) (Vega-Lugo et al., 2022). The RandC measure is determined as the colocalization extent when the target and reference are divided into subsets based on randomized condition (C) positions within the cell area (Vega-Lugo et al., 2022). Through conditional colocalization analysis, the extent of CD36 colocalization with active ITGB1 was assessed for the subset of CD36 associated with CD9 (*p(TwR|TwC)*), Fig. 3C) and the subset of active ITGB1 associated with CD9 (*p^rs^(Tw(RwC))*), Fig. 3C). This analysis revealed that CD36 colocalization with active ITGB1 was enhanced for the subset of ITGB1 associated with CD9, as for this subset (rightmost boxplot triplet in Fig. 3C) the extent of colocalization (*p^rs^(Tw(RwC))*) was significantly greater than its randomization counterparts (nullTR and RandC) and the overall extent of colocalization (p(TwR)). The converse was not true – the extent of CD36 colocalization with ITGB1 was not enhanced for the subset of CD36 associated with CD9 (middle boxplot triplet in Fig. 3C).

To facilitate the visualization of these conditional colocalization analysis results, we generated graphs indicating the extent of the colocalization with a circle (with the size reflecting the colocalization extent) and reporting the least significant p-value among the three comparisons (p(TwR) or nullTR or RandC) and expressed as a -log_10_(p-value) (values above 1.3 for 2 molecules and 1.7 for 3 molecules comparison indicate p<0.05) (Fig. 3D, E and F). These graphs illustrate that CD36’s colocalization with CD9 did not affect CD36’s colocalization with active ITGB1 (Fig. 3D) as CD36 colocalization with active ITGB1 when CD36 is with CD9 was not significantly different from the coincidental RandC measurements (Fig. 3C, middle) nor the overall colocalization of CD36 with active ITGB1 (Figure 3C and D). However, the colocalization of CD36 with ITGB1 when ITGB1 is with CD9 was significantly greater than the coincidental colocalization measures and the overall colocalization of CD36 with active ITGB1 (Figure 3C and D). Therefore, active ITGB1’s colocalization with CD9 significantly enhances CD36’s colocalization with active ITGB1.

Next, we investigated active ITGB1 as the condition, to determine the effect of ITGB1 on the colocalization between CD36 and CD9. To measure the effect of CD36 colocalization with active ITGB1 on the colocalization between CD36 and CD9, we determined the extent of CD36 colocalization with CD9 when CD36 is with active ITGB1 (middle, Fig. 3E) or when CD9 is with active ITGB1 (bottom, Fig. 3E). This analysis indicated that CD36’s colocalization with active ITGB1 does not enhance CD36’s colocalization with CD9, but, instead it is when CD9 is with active ITGB1 that the colocalization between CD36 and CD9 (∼13%) is significantly greater than the controls and overall colocalization of CD36 with CD9 (Fig. 3E). Therefore, we can conclude that CD9’s colocalization with ITGB1, enhances CD36’s colocalization with active ITGB1.

Conditional colocalization experiments performed between CD36, TfR and active ITGB1 as a negative control revealed that active ITGB1 had no influence on the colocalization between CD36 and TfR (Fig. 3F). All in all, from these conditional colocalization experiments, we observed that active ITGB1’s colocalization with CD9 enhances CD36 colocalization with active ITGB1, and that CD9’s colocalization with active ITGB1 enhances CD36’s colocalization with CD9. These conditional effects of CD9 and ITGB1 provide evidence for ternary complex formation between CD36, CD9 and ITGB1. To further investigate the organization of the complex, we evaluated CD36 interaction with CD9 or active ITGB1 via PLA at the surface of TIME cells silenced for ITGB1 or CD9, respectively, via stable expression of small hairpin RNAs (shRNA).

### Investigation of CD36-ITGB1-CD9 ternary complex via ITGB1 and CD9 silencing

We engineered TIME mEm-CD36 and TIME HT-CD36 cell lines to stably express shRNA targeting CD9 or ITGB1, using lentiviral transductions. Within these TIME cells, each shRNA significantly reduced the expression of ITGB1 or CD9 by ∼95% and ∼80%, respectively (Fig. 4A, B) (measured via immunoblotting, Suppl. Fig. 3).

**Figure 4:**
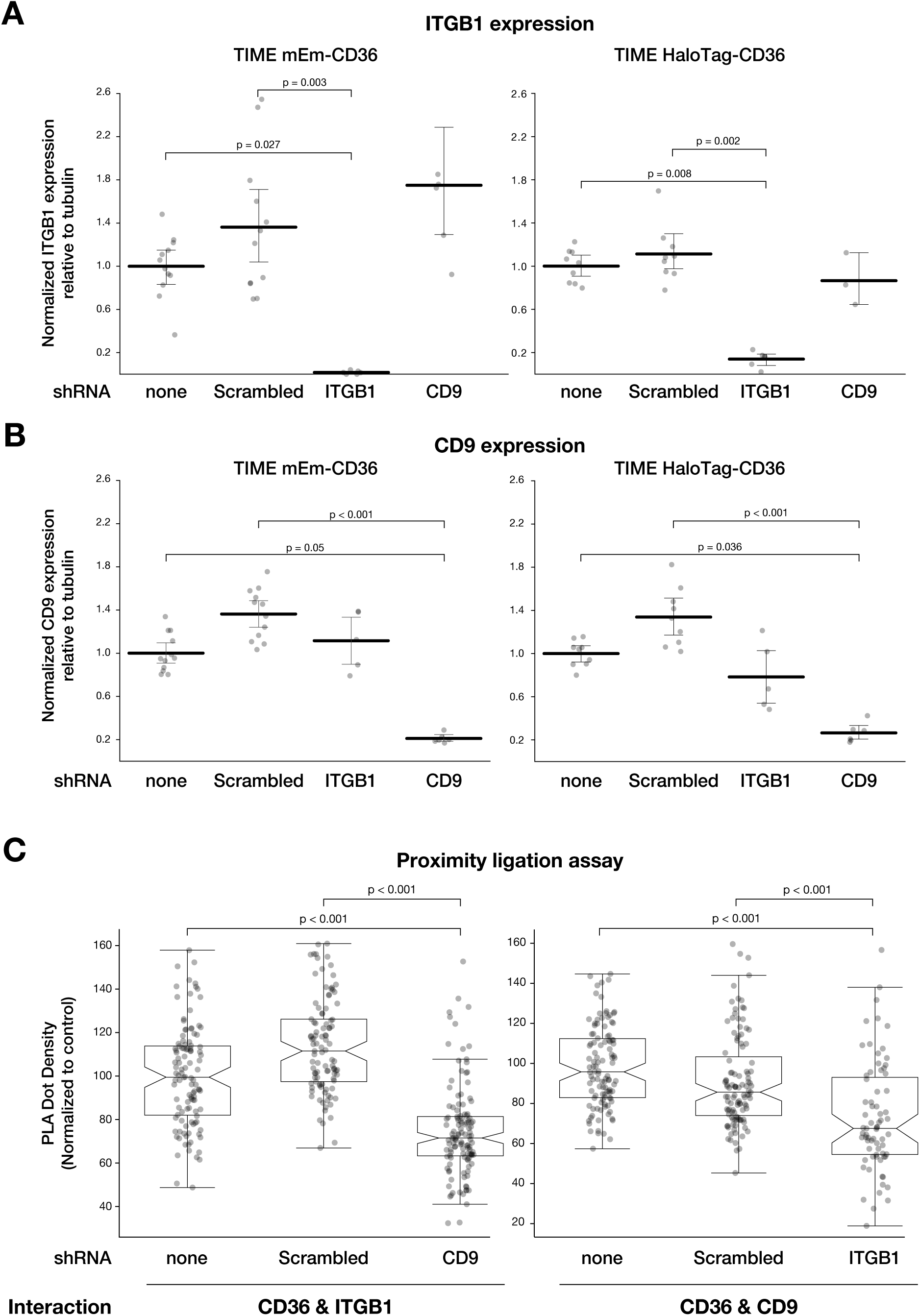
Silencing of CD9 and ITGB1 and PLA with CD36. (**A-B**) Levels of downregulation (**A**) of ITGB1 and (**B**) of CD9 in TIME mEm-CD36 and TIME HT-CD36 cells, as labeled on top of each graph. Whole cell lysates from stable cell lines expressing or not shRNAs indicated below the graphs were run on SDS-PAGE, transferred to nitrocellulose and immunoblotted for ITGB1 and CD9. Quantification of the protein levels relative to tubulin, were normalized to the expression level of cells not transduced with any shRNA. Statistical analysis was done using a non-parametric Kruskal-Wallis test followed by Dunn’s post-hoc test to determine pairwise significance values. Data from at least 3 independent experiments. (**C**) PLA measurements of CD36 interaction with active ITGB1 (left panel) and CD9 (right panel) in cells expression CD9 or ITGB1 shRNAs, respectively. PLA was performed as for Fig. 2B, on TIME mEm-CD36 expressing shRNAs as indicated on each graph. The boxplots indicate the PLA dots density normalized to measurement made in control cell lines, not expressing any shRNAs. Statistical analysis was done using a non-parametric Kruskal-Wallis test followed by Dunn’s post-hoc test to determine pairwise significance values. Data are from 3 experiments and a minimum of 100 cells analysed.

Through PLA experiments and quantifications, we determined that in TIME mEm-CD36 cells silenced for CD9, there was a significant (∼25%) reduction in the interaction between CD36 and ITGB1, shown by the lower density of PLA compared to control TIME mEm-CD36 (Fig. 4C). Similarly, ITGB1 knockdown (KD) affected the interaction between CD36 and CD9, with a reduction of the density of PLA dots by ∼18% compared to TIME mEm-CD36 cells (Fig. 4C). Overall, the PLA analysis revealed that CD36 interaction with ITGB1 is positively influenced by CD9 and vice versa. In addition, the inactivation of CD9 has a larger effect on CD36’s interaction with ITGB1 than ITGB1 KD has on the interaction between CD36 and CD9. These findings align with our conditional colocalization results, showing that ITGB1 colocalization with CD9 has a greater effect on CD36’s colocalization with active ITGB1 than CD9 colocalization with ITGB1 has on CD36’s colocalization with CD9. This suggests that CD9 is an important molecule to connect CD36 to active ITGB1. PLA and conditional colocalization experiments and specific silencing of ITGB1 or CD9 provide support for this ternary complex forming between CD36, ITGB1 and CD9. We next designed experiments to evaluate the role of CD9 and ITGB1 in the context of CD36 signaling.

### Examining the role of CD9 and active ITGB1 in CD36-Fyn signaling

To investigate whether the CD36-ITGB1-CD9 complex plays a role in signal transduction to activate Fyn, we determined levels of the active form of the Src family kinase (SFK) using an antibody directed against the phosphorylated tyrosine 420, conversed among SFK (pSFK). While the phosphorylation of other SFK would possibly be detected with this antibody, several lines of evidence support the fact that the main SFK involved downstream of CD36 is Fyn. First, in HMVEC cells, we have previously shown, through specific siRNA silencing, that the main contributor of pSFK signal downstream of CD36 was Fyn (Githaka et al., 2016). Second, Fyn is highly expressed in TIME cells (data from Human Protein Atlas, https://www.proteinatlas.org/), and third, Fyn was found to be significantly enriched in our BAR proximity biotinylation experiment, while other SFK were not (Fig. 1D).

To specifically engage CD36 and enhance its clustering, a multivalent IgM mouse anti-CD36 antibody was used. This multivalent ligand is a known activator (Dawson et al., 1997; Jiménez et al., 2000a; Githaka et al., 2016) that has no confounding effect that the natural ligand TSP-1 could have by also binding to other receptors, including ITGB1 (Calzada et al., 2004)e. Following IgM stimulation, TIME HT-CD36 cells were stained with rabbit anti P-Y420-Src and imaged by TIRF microscopy. Mean intensity of pSFK was determined, showing that IgM stimulation resulted in a ∼ 1.7-fold increase in pSFK levels compared to unstimulated cells (Fig. 5A). Similarly, TIME HT-CD36 cells expressing scrambled shRNA exhibited a ∼1.54-fold increased pSFK signal upon stimulation (Fig. 5A). The significant increase in pSFK intensity following stimulation for both cell lines confirms that multivalent stimulation of CD36 at the surface of TIME-HT CD36 cells results in Fyn activation (Githaka et al. 2016). However, stimulation of TIME HT-CD36 cells expressing ITGB1 shRNA resulted in no change in pSFK mean intensity compared to unstimulated control (Fig. 5A). Stimulation of CD9 shRNA knockdown cell lines resulted in ∼1.3-fold change in pSFK signal (Fig. 5A) which is significantly higher than the unstimulated control but attenuated in comparison to the response seen in the wildtype TIME HT-CD36 cells. Therefore, we concluded that CD36’s capacity to activate Fyn is abolished in the absence of ITGB1. Next, we explored by conditional colocalization analysis the relationship between CD36, CD9, ITGB1, and pSFK during stimulation.

**Figure 5:**
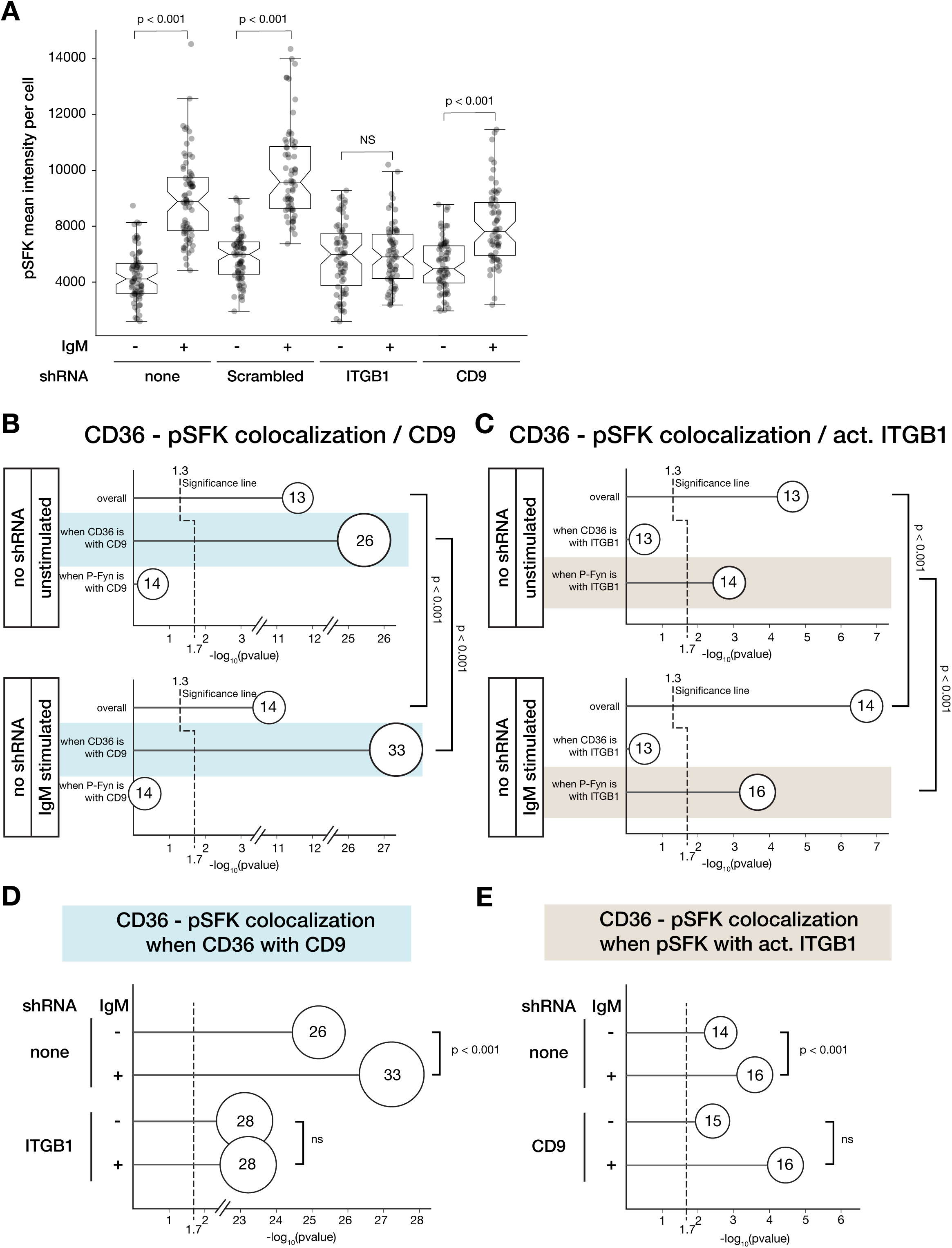
Role of ITGB1 and CD9 in CD36-Fyn signaling. (**A**) Levels of active Fyn determined by immunostaining of pSFK in cells stimulated or not with anti-CD36 IgM antibodies. TIME HT-CD36 cells expressing no shRNAs or scrambled, CD9 or ITGB1 shRNAs were stimulated (+) or not (-) with mouse anti-CD36 IgM (10 µg/mL) for 15 min. Cells were fixed, immunostained for pSFK and imaged by TIRF microscopy using identical parameters for all conditions. The levels of pSFK were determined using Cell Profiler by measurement of the mean intensity of pSFK for each cell. Cell segmentation was done using F-actin labeling. The boxplots indicate the median intensity of pSFK per cell and statistical comparisons were done using non-parametric T-test between stimulated and unstimulated conditions. Results are from 3 experiments and a minimum of 64 cells analyzed per condition. (**B**) Summary of conditional colocalization measurements indicating the effect CD9 on the colocalization between CD36 and pSFK. TIME HT-CD36 cells were stimulated (bottom panel) or not (top panel) with anti-CD36 IgM for 15 min, and immunostained for conditional colocalization with HT-JF549X, anti-pSFK-AF488 and anti-CD9 AF647. Statistical analysis as for Fig. 3D, E, and F. A minimum of 75 cells per conditions were imaged across 3 experiments. (**C**) Summary of conditional colocalization measurements indicating the effect of active ITGB1 on the colocalization between CD36 and pSFK as in (B). TIME HT-CD36 cells were stimulated (bottom panel) or not (top panel) with anti-CD36 IgM for 15 min, and immunostained for conditional colocalization with HT-JF549X, anti-pSFK-AF488 and anti-active ITGB1 AF647. At least 53 cells per condition were imaged across 3 experiments. Comparisons shown as in (C). (**D-E**) Effect of silencing ITGB1 (**D**) or CD9 (**E**) on CD36 and pSFK conditional colocalization measurements highlighted in panel B (blue) and C (brown). TIME HT-CD36 cells expressing indicated shRNA and stimulated or not with IgM anti-CD36 were processed for conditional colocalization between CD36, pSFK and either CD9 (D) or ITGB1 (E). A non-parametric Kruskal-Wallis test followed by Dunn’s post-hoc test were used to determine statistical differences between stimulated and unstimulated conditional colocalization experiments. Data are from 3 experiments and a minimum of 50 cells analysed.

### Role of ITGB1 and CD9 in CD36-Fyn signalling

To determine the contribution of CD9 and ITGB1 to CD36 complexes that promote Fyn activation, we performed conditional colocalization on TIME HT-CD36 stimulated with mouse anti-CD36 IgM and measured the influence of CD9 and active ITGB1 on the colocalization of CD36 with pSFK (Fig. 5B and C). At steady-state, ∼13% of CD36 molecules colocalized with pSFK (Fig. 5B and C). Upon IgM stimulation, there was a significant increase in the overall CD36 colocalization with pSFK (Fig. 5B and C).

When CD9 or active ITGB1 were used as condition (C) for the analysis, they were found to positively influence the CD36-pSFK colocalization. At rest, the analysis indicated that CD36-pSFK was found where pSFK colocalized with ITGB1 (Fig. 5C, highlighted in brown) and even more significantly when CD36 colocalizes with CD9 (Fig. 5B, highlighted in blue). When stimulated with anti-CD36 IgM, both populations increased significantly compared to unstimulated (Fig. 5B and C), indicating that productive ternary complexes are those where CD36 colocalizes with CD9 and where ITGB1 is associated with pSFK.

Next, similar conditional colocalization analyses were performed in cells silenced for CD9 or ITGB1. Fig. 5D and E focus on the changes occurring in the highlighted populations from panels B and C. Inactivation of either protein resulted in a loss of the increased colocalization extent seen for CD36-pSFK compared to control condition. These data reinforce the critical role of both ITGB1 and CD9 in creating productive ternary complexes with CD36.

Conditional colocalization analysis revealed that CD9 enhanced CD36’s colocalization with pSFK, as CD36-CD9 colocalization events had a ∼2-fold increase colocalization in comparison to overall CD36-pSFK colocalization (Fig. 5B). IgM stimulation resulted in a significant increase in this subpopulation from ∼26% at rest to 33% (Fig. 5B). The higher proportion of CD36-CD9 colocalization events associated with pSFK upon CD36 IgM stimulation was lost when ITGB1 was silenced (Fig. 5D). Together with the elimination of pSFK stimulation in cells expressing ITGB1 shRNAs (Fig. 5A), our results indicate that in the absence of ITGB1, the coupling between CD36 and CD9 still exists (albeit reduced; Fig. 4C), but that IgM clustering of CD36 in these complexes does not result in Fyn activation.

Conditional colocalization analysis also revealed changes in CD36’s interaction with ITGB1 and pSFK in response to CD36 stimulation. Contrary to the effect of CD9 (whose association with CD36 increases CD36 colocalization with pSFK; Fig. 5B), we found that it is not the interaction of CD36 with active ITGB1 that favors CD36 connection to pSFK, but more the colocalization of active ITGB1 with pSFK that positively influence CD36-pSFK colocalization (Fig. 5C). Upon IgM stimulation, the proportion of pSFK-active ITGB1 colocalization events also colocalizing with CD36 (Fig. 5C, highlighted in brown) significantly increase from ∼14% to ∼16%. This suggests that active ITGB1 has a more direct role in the activation of Fyn, as its connection with pSFK has a more significant effect than that of CD36. Within the CD9 KD cell line, the IgM-induced increase in this population of complexes (where CD36 colocalizes with pSFK when pSFK is with ITGB1) was abolished (Fig. 5E). Altogether, these results demonstrate that CD9 has a critical role in associating CD36 to complexes containing active ITGB1 sites to promote Fyn activation.

## Discussion

### Identification of CD36 interacting partners with BAR and PLA methods

By adapting a proximity labeling approach initially described by Bar et al. (Bar et al., 2018) we targeted the scavenger receptor CD36 at the surface of endothelial cells to reveal a set of proteins potentially implicated in the TSP-1 mediated anti-angiogenic response. This method has the advantage to enable interrogation of the composition of molecular complexes in a somewhat preserved cellular environment, and it represent a powerful alternative to co-immunoprecitation for which solubilization of proteins could often result in a loss of indirect, transient, or weak interactions. Known interactors of CD36, described in different cellular contexts were confirmed. Using proximity ligation assays, we confirmed the proximity of 11 candidate proteins (CD59, CD9, ITGA5, PTP1B, CD146, CD146, ITGA2, ITGB1, MYOF, CD151, and CD81) to CD36 in microvascular endothelial cells within a ∼40 nm radius of CD36. To further validate these interactions, we proceeded to investigate the roles of CD9 and ITGB1 in CD36-induced anti-angiogenic signaling and organization.

### Model of CD36-Fyn signal transduction

Our data support a model in which CD36, CD9 and ITGB1 form a ternary complex, in which CD9 plays a bridging role between the scavenger receptor and the integrin. In this multimolecular assemblies, active ITGB1 appears critical to promoting Fyn activation. We found that IgM stimulation resulted in a significant increase in both the fraction of CD36 with pSFK when CD36 is with CD9 or when active ITGB1 is with pSFK. Given that these interactions are already in place at steady-state and become more prevalent upon IgM stimulation, it suggests that enhancement of CD36 clusters lead to the formation and strengthening of these productive complexes towards increasing levels pSFK.

### CD9 as a bridge between CD36 and ITGB1

The tetraspanin CD9 appears to play a critical role in bridging CD36 with ‘productive’ ITGB1 complexes. This was demonstrated by the facts that CD36 and ITGB1 colocalization was positively influenced by their colocalization with CD9 (Fig. 3D and E) and that downregulation of CD9 resulted in a significant decrease in CD36-ITGB1 interaction (Fig. 4C). Furthermore, CD36 colocalization with pSFK, at rest and following stimulation, was strongly correlated to CD36 being with CD9, indicating that complexes containing CD36-CD39 and ITGB1 are supportive of Fyn activation. However, when CD9 was downregulated, there was only a partial reduction in the IgM induced increase in pSFK levels (Fig. 5A). This might be the result of residual interactions between CD36 and ITGB1 in cells silenced for CD9 (Fig. 4C), which could be supported by compensatory tetraspanins, such as CD151 or CD81 that we have found to also interact with CD36 (Fig. 2B).

Studies in macrophages have shown that CD36’ s interaction with CD9 is mediated by the GXXXG motif within the N-terminal transmembrane domain of CD36 (Huang et al., 2023; Jana et al., 2025). Since CD9 also contains a GXXXG motif in the 3^rd^ transmembrane domain, we hypothesized that GXXXG motifs mediate the transmembrane domain interaction (Teese and Langosch, 2015; Huang et al., 2023). In fact, the mutation G12V in CD36 resulted in a drastic loss in its capacity to colocalize with ITGB1 and CD9 (Jana et al., 2025). Tetraspanin CD81 may facilitate CD36’s interaction with ITGB1 in the absence of CD9, as it also contains a transmembrane GXXXG motif and has been shown to associate with CD36 (Heit et al., 2013) and ITGB1 (Serru et al., 1999; Oguri et al., 2020). Although CD151 does not have a transmembrane GXXXG motif, studies have revealed that CD151 associates with integrin α3β1 (Yauch et al., 1998, 2000; Berditchevski et al., 2002; Zhang et al., 2002) and CD36 (Kazerounian et al., 2011). Therefore, it is assumed that CD36 forms a complex where CD151 connects CD36 to α3β1 (Thorne et al., 2000; Berditchevski et al., 2002). Our PLA association analysis confirmed interaction between CD36-CD81, and CD36-CD151, however, to lower levels than CD36 with CD9. Conditional colocalization experiments between CD36-ITGB1 and CD81 or CD151 would help determine whether CD81 and/or CD151 compensate for the absence of CD9. Comparing CD36 anti-angiogenic signaling within TIME cells with all three tetraspanins CD9, CD81 and CD151 knocked down would allow us to determine their role in CD36-Fyn signaling. However, our data strongly support the notion that CD9 plays a major role in facilitating interactions needed for CD36 multimolecular complex organization and signaling.

In addition, and like the effect of tetraspanins on epidermal growth factor receptor (EGFR) (Sugiyama et al., 2023), association of CD36 with CD9 could potentially be modulating its entrapment within nanodomains in a specific and more favorable conformation to interact with active ITGB1. Further analysis of the mobility of CD36 in cells lacking CD9 and other tetraspanins would be necessary to uncover this possibility.

### ITGB1 is critical for Fyn activation

If CD9 appears to connect CD36 to active ITGB1, what is the function of ITGB1 in this complex? ITGB1 has been shown to be essential for Fyn activation in other biological processes such as oligodendrocyte stimulation with Aβ oligomers (Quintela-López et al., 2019), Schwann cell adhesion and differentiation (Chen et al., 2000b), integrin-signaling (Wary et al., 1998), and oligodendrocyte differentiation (Liang et al., 2004). Moreover, CD36’s interaction with ITGB1 was shown to have a functional role in macrophages, as the knockdown of ITGB1 inhibited Src and spleen tyrosine kinase activation upon CD36-mediated oxLDL uptake (Heit et al., 2013). CD36 interaction with integrin α6β1 was identified within microglial cells (Koenigsknecht and Landreth, 2004; Bamberger et al., 2003). Antibody blocking of CD36 or ITGB1 inhibited ß-amyloid-mediated activation of Fyn and intracellular tyrosine kinase signaling (Bamberger et al., 2003). Here, we show how ITGB1 is needed for CD36-Fyn anti-angiogenic signalling in microvascular ECs. Although this current study reveals ITGB1 to be necessary for signaling, the specific mechanism by which ITGB1 promotes Fyn activation is still enigmatic. One example where integrins were linked to activation of Src-family kinases is during Fcγ-mediated phagocytosis (Freeman et al., 2016). Similar to this earlier study, ITGB1 could be facilitating CD36-Fyn signaling by creating an area in which inhibitory phosphatases would be excluded, tipping the balance towards more Fyn trans-autophosphorylation and activation. In this context, the connection with the actin cortex for promoting these exclusion zones would be important and could explain dependencies already established between CD36, the actin cortex and Fyn activation (Githaka et al., 2016; Dasgupta et al., 2023).

Integrins are heterodimeric proteins, and while we have examined the specific role of ITGB1, we have still to define whether specific integrin alpha/beta heterodimers are required for CD36-induced signaling. In our BAR proximity biotinylation experiments, we have identified that integrin alpha-2, -3, -5, and -6 are enriched in proximity to CD36. PLA interaction analysis for these alpha subunits showed that only alpha-5 and alpha-2 interact with CD36 (Fig. 2B). Further conditional colocalization analysis between CD36, ITGB1, and integrin-alpha 2, and -5, would be required to elucidate which integrin-alpha/ITGB1 heterodimer is implicated in CD36 signaling.

## Conclusion

Through employing proximity labeling (BAR) and ligation (PLA) we have discovered the identity of proteins within CD36 multimolecular complexes. Of the candidate proteins identified, we revealed the roles of CD9 and ITGB1 in CD36 signaling. Silencing and conditional colocalization experiments investigating the interaction between CD36, CD9, and active ITGB1 provided support for these proteins to form a ternary complex, in which CD9 enables CD36 to contact ITGB1, the latter playing a critical role in downstream signaling and activation of Fyn. We have demonstrated how ITGB1 and CD9 play a crucial role in facilitating CD36 organization and function. This investigation of CD36 has provided a roadmap that can be applied to study the organization of many membrane proteins and better understand how multimolecular complexes provide modularity to promote specificity and signal amplification. Additionally, the roles of lipid nanodomains and the actin cortex in organizing these platforms should also be considered and warrant further investigation.

## Materials and Methods

### Cell Culture

Human telomerase (hTERT)-immortalized microvascular endothelial cells isolated from human foreskin (TIME, ATCCⓇ CRL-4025) were cultured in Vascular Cell Basal Medium (ATCCⓇ PCS-100-030) supplemented with Microvascular Endothelial Cell Growth Kit-VEGF (ATCCⓇ PCS-110-041), 0.5 mg/mL Penicillin/Streptomycin and 12.5 µg/mL blasticidine (ThermoFisher Scientific, Waltman, MA, USA, ref: A11139-03) and grown at 37°C in 5% CO_2_ atmosphere.

Lentiviral transduction was used to generate stable TIME cell lines expressing HaloTag-CD36 (HT-CD36) or mEmerald-CD36 (mEm-CD36) under the control of a tetracycline-inducible promoter. HT-CD36 and mEm-CD36 were first cloned into pLVX-TRE3G vector (Takara Bio USA Inc., San Jose, USA) using AgeI and EcoRI restriction sites. HEK293T cells were transfected in parallel using LentiX packaging kits (Takara Bio, ref:631275) and pLVX-mEmCD36 or pLVX-HT-CD36 (transfer vector with mEm-CD36 or HT-CD36) and pLVX-Tet3G (transfer plasmid containing the tetracycline-dependent transactivator, rtTA). Media containing viral particles were harvested after 48h, filtered (0.45µm pore size) and mixed with 10 µg/mL polybrene. To generate the tetracycline-inducible TIME cell lines, both pLVX-Tet3G and either pLVX-mEmCD36 or pLVX-HT-CD36 viral supernatants were added to TIME cells at 50% confluency and incubated for 72h before applying 0.5 µg/mL of G418 (resistance gene contained in pLVX-Tet3G) and 2 µg/mL of puromycin (resistance gene for mEm/HT-CD36). After two weeks of selection, surviving cells were collected and plated in parallel on coverslips to evaluate the inducible expression of mEmerald-CD36.

Lentiviral transduction was also used to generate TIME HT-CD36 and TIME mEm-CD36 cells for stable expression of ITGB1, CD9 or scrambled shRNAs. Plasmids expressing scrambled, ITGB1, or CD9 shRNA were obtained from VectorBuilder (VectorBuilder Inc., IL, USA) and consisted of a pLV vector, containing an hygromycin resistance cassette, and the shRNA under the control of a constitutively active U6 promoter. HEK293T cells were transfected in parallel using LentiX packaging kit (Takara Bio) and pLV-scrambled/CD9/ITGB1-shRNA transfer vectors. Media containing viral particles were harvested after 48h, filtered (0.45µm pore size), and mixed with 10 µg/mL polybrene. To generate the scrambled shRNA, CD9 shRNA, or ITGB1 shRNA KD cell lines, TIME HT-CD36, and TIME mEm-CD36 were transduced at 50% confluency for 48 hours. Following the transduction period, cells were incubated with TIME cell media supplemented with 0.2 mg/mL hygromycin (resistance gene for scrambled/CD9/ITGB1-shRNA). After 1 week of selection, surviving cells were collected and their lysate was analyzed via immunoblotting to confirm candidate protein down regulation.

### CD36 Biotinylation by Antibody Recognition (BAR)

TIME mEm-CD36 cells were grown to ∼80-90% confluency in 6 cm tissue culture dishes or on coverslips and induced with 0.5 µg/mL of doxycycline for 18 hours to allow mEm-CD36 expression. Cells were fixed with 4% PFA (Electron Microscopy Sciences, Hatfield, PA, USA) for 20 minutes on ice. Following fixation, 1 mL of 3% BSA in PBS was added to the cells for 1 hour at room temperature. Subsequently, 1mL of 1:1000 mouse anti-CD36 (clone 131.2, generous gift from Dr. Tandon, Cellphire, Rockville, USA) was added for 1 hour at room temperature. Cells were washed three times with PBS. 1 mL of Goat anti-mouse fragment antigen binding protein (Fab) conjugated to horseradish peroxidase (HRP) (Rockland, Baltimore, MD, USA) was diluted 1:1000 and incubated with the cells for 1 hour at room temperature. Following three washes with PBS, Biotin-XX-tyramide (Biotium, San Franscisco, CA, USA) diluted 1:1000 in the tyramide amplification buffer (Biotium, San Franscisco, CA, USA) was added to cells for 10 minutes at room temperature. Cells were protected from light, as the tyramide reaction is light sensitive. Following the reaction, cells were washed three times with PBS.

For analysis of cell lysates via immunoblot, 500 µL of lysis buffer (150 mM NaCl, 50 mM Tris-HCl at pH 7.2, 0.2% SDS, 1% Triton X-100, protease inhibitor cocktail) was added for 30 min on ice. Cell homogenates were harvested using a cell scrapper and centrifuged at 14,000 RPM for 20 minutes at 4°C. A fraction of the supernatant was kept as whole-cell lysate (WCL); the remaining supernatant was processed for streptavidin capture.

For analysis via fluorescence microscopy, coverslips were incubated for 1 hour with streptavidin coupled to Alexa Fluor 647 (Jackson Immunoresearch, Philadelphia, PA, USA) and donkey-anti mouse Cy3 (Jackson Immunoresearch, Philadelphia, PA, USA) diluted 1:500 in 3% BSA in PBS. Subsequently, cells were washed three times in PBS and post-fixed with 4% PFA (Electron Microscopy Sciences, Hatfield, PA, USA) for 10 minutes at room temperature. Cells treated with CD36 BAR were imaged via widefield microscopy.

### Streptavidin Capture and Tandem Mass Spectrometry

CD36 BAR lysates, Pierce Streptavidin Magnetic Beads (ThermoFisher, Waltman, MA, USA) and lysis buffer were added to a deep 96 well plate (ThermoFisher, Waltman, MA, USA). The KingFisher Duo Prime Purification System was used for automated capture. First, streptavidin beads and CD36 BAR lysates were incubated for 18 hours at 4°C. Following streptavidin capture, beads were successively transferred to wells containing 100 mM NH_4_CO_3_ for wash, then 10 mM dithiothreitol (DTT) for reduction, 50 mM iodoacetamide for alkylation, and 100 mM DTT for quenching, and finally proteins were digested with trypsin directly from streptavidin beads for 8 hours.

Trypsinized peptides were transferred to tubes and concentrated using speed-vacuum evaporation. Peptides were desalted using Thermo Scientific Pierce C18 Spin tips following the manufacturer’s instructions (ThermoFisher, Waltman, MA, USA). Peptides were resuspended in Buffer A (3.9% acetonitrile, 0.1% formic acid in mass spectrometry grade H_2_O) and the samples were analyzed on a nanoflow-LC (Thermo Scientific EASY-nLC 1200 system) coupled to the Orbitrap Fusion Lumos Tribrid Mass Spectrometer (Thermo Fisher Scientific). The reverse separation of the peptides was performed on a 15 cm analytical column (2 μm, 100 Å, 50 μm × 15 cm, PepMap RSLC C18; Thermo Fisher Scientific) with a 60-minute linear gradient from 3.9% to 36.8% acetonitrile in 0.1% formic acid. Raw data was processed using Proteome Discoverer (v2.4) against the Uniprot human proteome data (2020, 42285 sequences), trypsin was defined as the digestion, with a precursor mass tolerance of 15 ppm and 0.8 Da for fragment tolerance. For protein abundance calculation, the “LFQ and Precursor quant” workflow was applied with no normalization and no scaling of the data. Output from MS/MS includes peptide spectral matched (PSM), number of peptides, and protein intensity. Protein intensity is the sum of the signal intensity for all the peptides corresponding to an identified protein (Vogel and Marcotte, 2009).

### Normalization of Protein Intensities for CD36 BAR MS/MS Dataset

MS/MS identified ∼800 for CD36 BAR experiments (TIME mEm-CD36 induced with doxycycline for mEm-CD36 expression, n=8) and ∼1000 proteins for BAR control (TIME mEm-CD36 not induced, n=8) (Table 1). The number of proteins identified was equivalent for all the replicates within each treatment (n=8). However, there was a large variation in average protein intensity between replicates. To nullify the variance and allow for comparisons between the 16 samples (8 for CD36 BAR and 8 for BAR control), we developed an intensity normalization method.

Normalization was performed by finding the common proteins to all CD36 BAR (n=8) and BAR control (n=8) replicates (illustrated on Suppl Fig. 2A). 367 commonly identified proteins were utilized to determine the normalization factor to be applied to each of the 16 samples. Prior to applying normalization, the MS data was zero-filled by replacing blank protein intensity values with the minimum protein intensity identified within each sample (Suppl Fig. 2B). For each replicate, the average protein intensity of commonly identified proteins was determined. The normalization factor was then calculated by dividing the average common protein intensity for the replicate by the lowest average of common protein intensity across all experiments. CD36 BAR replicate 8 had the lowest average common protein intensity and its average was used to calculate the normalization factor. The normalization factor for each replicate was calculated as the average common protein intensity within the replicate divided by the one of CD36 BAR replicate 8 (Suppl Fig. 2A).

Normalization effectively reduced variations between all replicates to an average protein intensity of ∼0.5×10^6^ (Suppl Fig. 2B). Boxplots of the protein intensities for CD36 further demonstrate the effectiveness of this method, as the interquartile ranges are greatly reduced after normalization (Suppl. Fig. 2C). Normalization of protein intensities reduced variance in the protein intensities among samples allowing us to make statistical comparisons between CD36 BAR and CD36 BAR control data. Candidate proteins were defined as those proteins with a significantly higher intensity in the CD36 BAR condition (determined via Welch’s T-test), and with an enrichment ratio (intensity in CD36 BAR / intensity on BAR control) higher than 2. This list contains 164 candidate proteins and is attached as supplementary table 1.

### Immunoblotting

Whole cell lysates from BAR experiments or from shRNA knockdown expression analysis were resolved on 10% SDS-PAGE for 1 hour at 150 V. Proteins were transferred to nitrocellulose membranes for 1 hour at 110 V which were then blocked using 3% BSA dissolved in Tris Buffered Saline Tween (TBST) at room temperature for 45 minutes. Membranes were incubated overnight with labeled streptavidin or primary antibody solution at 4°C as indicated on Table 2. Following primary antibody incubation, membranes were washed three times with TBST and then incubated with secondary antibody solutions for 1 hour at room temperature (Table 2). Membranes were visualized using Li-Cor Odyssey (Li-Cor Bioscience, Lincoln, NE, USA). Following visualization, Empira Studio Software (Li-Cor Bioscience, Lincoln, NE, USA) was used to quantify protein knockdown for our TIME mEm/HT-CD36 expressing scrambled, ITGB1 or CD9 shRNA cell lines. Relative expression of ITGB1 or CD9 was normalized to housekeeping protein control, tubulin.

**Table 2:**
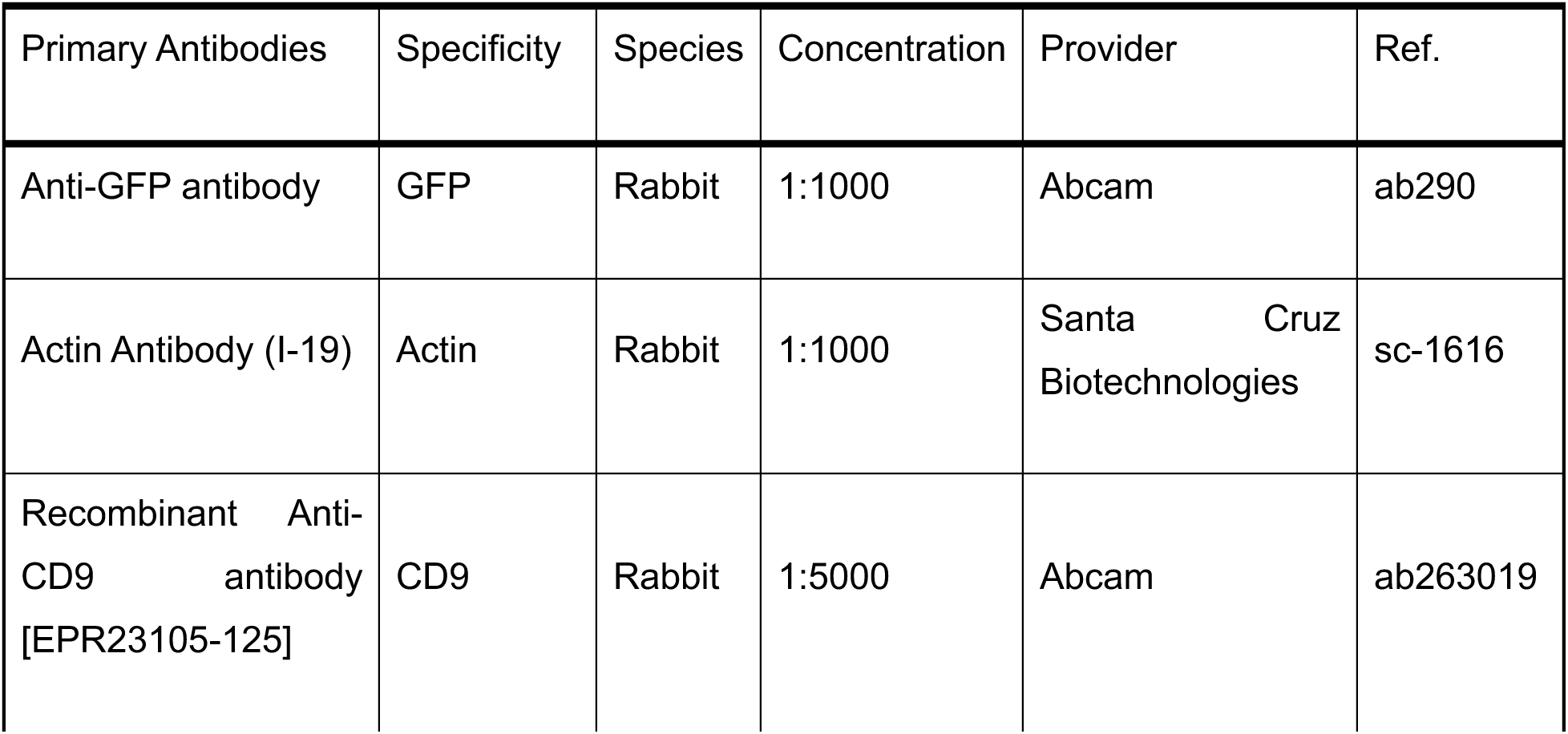
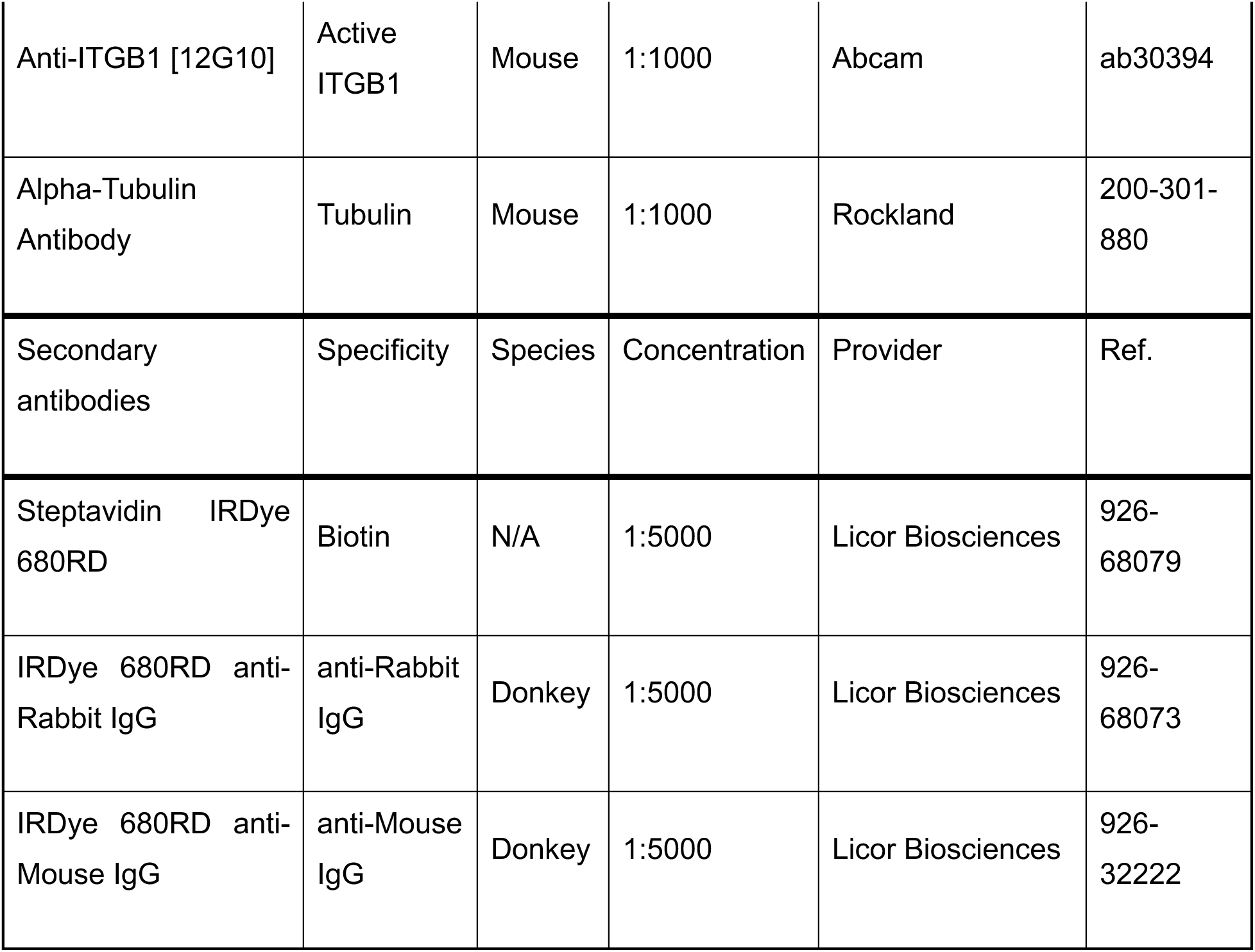
List of primary and secondary antibodies for immunoblot experiments.

### Duolink PLA Protocol

TIME mEmCD36 cells were plated on 18 mm glass coverslips (Electron Microscopy Sciences, Hatfield, PA, USA, ref: 72290-08) and induced with 0.5 µg/mL of doxycycline. Next day, cells were fixed using 4% paraformaldehyde (Electron Microscopy Sciences, Hatfield, PA, USA) for 20 minutes on ice and washed three times with PBS. Cell surface proteins were labeled for 30 min with 10 µg/mL of lectin-binding protein concanavalinA coupled to Alexa Fluor (AF)-488 (ThermoFisher, Waltman, MA, USA) diluted in 3% BSA in PBS with the purpose of providing a fluorescence signal to allow segmentation of the cells area for quantification of PLA images (Kiernan, 1975).

Cells were then washed three times with PBS to remove unbound concanavalinA-AF488, permeabilized with PBS containing 3% BSA + 0.1% Triton X-100 (Avantor, Radnor, PN, USA) for 30 minutes at room temperature. Cells were subsequently washed with PBS and two drops of Duolink blocking solution (1% BSA) (Millipore Sigma, Burlington, MA, USA) were added to each coverslip for 45 min at 37°C. Primary antibodies diluted in the Duolink blocking solution were added to each coverslip for 1 hour at room temperature. Antibodies pairs used for PLA are listed on Table 3. Coverslips were washed two times with Wash Buffer A (50 mM NaCl, Tris-base 10mM, tween 0.05%, pH 7.4) for 5 minutes each at room temperature. PLA donkey anti-rabbit PLUS and donkey anti-mouse MINUS probes (Millipore Sigma, Burlington, MA, USA) were diluted 1:5 in Duolink blocking solution, and 40 µL was added to coverslips for 1 hour at 37°C. Next, cells were washed twice with buffer A for 5 minutes each at room temperature. For probe ligation, Ligation Stock Buffer (Millipore Sigma, Burlington, MA, USA) was diluted 1:5 in double distilled water, and Ligase from the PLA kit (Millipore Sigma, Burlington, MA, USA) was added at a 1:40 dilution in ligase buffer. 40 µL of the diluted ligase solution was added to each coverslip and incubated at 37°C for 30 minutes. Following this incubation, coverslips were washed twice with Wash Buffer A for 5 minutes at room temperature. For amplification, Amplification Stock Buffer (Millipore Sigma, Burlington, MA, USA) was diluted 1:5 in double distilled water, and Polymerase from the PLA kit was added 1:80 to the diluted amplification solution. 40 µL of the diluted amplification buffer was added to each coverslip and incubated at 37°C for 100 minutes. Coverslips were washed twice with Wash Buffer B (100 mM NaCl, 35 mM Tris-base, Tris-HCl 215 mM, pH 7.4) for 10 minutes at room temperature. For estimation of cell number, 4′,6-Diamidino-2-phenylindole dihydrochloride, 2-(4-Amidinophenyl)-6-indolecarbamidine (DAPI) (Millipore Sigma, Burlington, MA, USA) diluted 1:500 in 3% BSA-PBS was incubated with coverslips for 30 minutes at room temperature. Coverslips were quickly rinsed twice with Wash Buffer B and then post-fixed using 4% PFA at room temperature for 10 minutes. Finally, cells were washed with 1:100 Wash Buffer B for 1 minute and mounted on slides with Prolong Glass Antifade mounting media (ThermoFisher, Waltman, MA, USA).

**Table 3:** List of primary antibodies used in PLA experiments.

| Primary Antibodies | Specificity | Species | Concentration | Provider | Catalog No. |
| --- | --- | --- | --- | --- | --- |
| Purified anti-human CD9 Antibody | CD9 | Mouse | 1:1000 | Biolegend | 312102 |
| Anti-CD36 (Clone131.2) | CD36 | Mouse | 1:1000 | Gift from Dr. Tandon | N/A |
| Anti-human CD59 | CD59 | Mouse | 1:1000 | Biolegend | 304702 |
| Anti-CD146 antibody | CD146 | Mouse | 1:1000 | Abcam | ab24577 |
| Anti-CD151 antibody | CD151 | Mouse | 1:500 | Abcam | ab33315 |
| Anti-GFP antibody | GFP | Rabbit | 1:1000 | Abcam | ab290 |
| Anti-GFP antibody | GFP | Mouse | 1:1000 | Abcam | ab1218 |
| Human EPCR Antibody | EPCR | Mouse | 1:200 | R&D Systems | MAB22451-SP |
| Anti-Integrin beta-1 antibody [12G10] | Active ITGB1 | Mouse | 1:1000 | Abcam | ab30394 |
| Anti-Integrin alpha 2 | ITGA2 | Rabbit | 1:1000 | Abcam | ab209766 |
| Anti-Integrin alpha 3 antibody | ITGA3 | Rabbit | 1:1000 | Abcam | ab131055 |
| Anti-Integrin alpha 5 | ITGA5 | Rabbit | 1:1000 | Abcam | ab150361 |
| Anti-Mitofilin antibody [2E4AD5] | Mitofilin | Mouse | 1:1000 | Abcam | ab110329 |
| Recombinant Anti-Myoferlin antibody [EPR18887] | Myoferlin | Rabbit | 1:200 | Abcam | ab178386 |
| Anti-PTP1B antibody [4F8F11] | PTP1B | Mouse | 1:100 | Abcam | ab201974 |
| Anti-Stomatin antibody | Stomatin | Rabbit | 1:200 | Abcam | ab169524 |

**Table 4:**
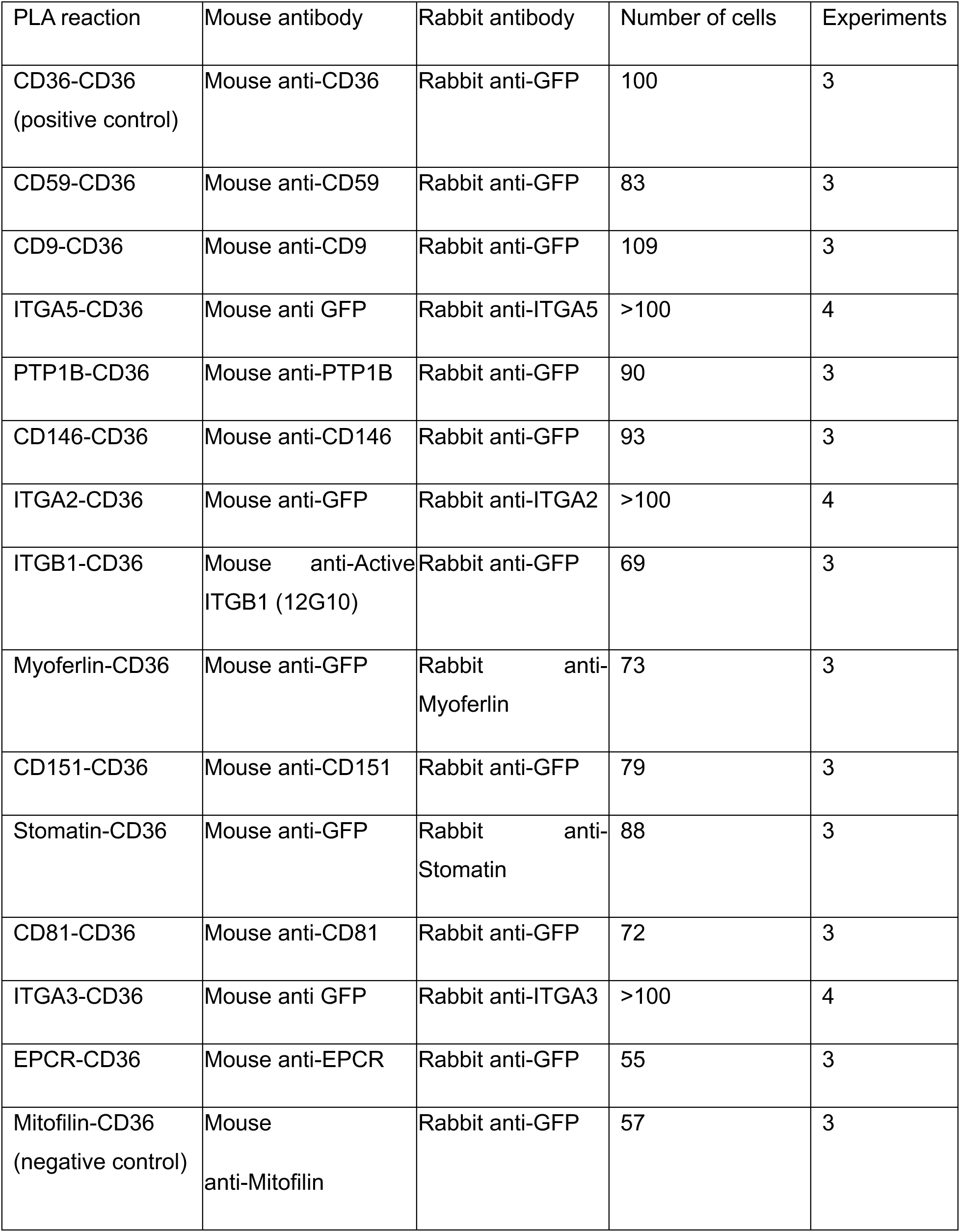
Antibody pairs, technical replicates and final number of cells imaged in PLA reactions between CD36 and candidate proteins.

| PLA reaction | Mouse antibody | Rabbit antibody | Number of cells | Experiments |
| --- | --- | --- | --- | --- |
| CD36-CD36<br>(positive control) | Mouse anti-CD36 | Rabbit anti-GFP | 100 | 3 |
| CD59-CD36 | Mouse anti-CD59 | Rabbit anti-GFP | 83 | 3 |
| CD9-CD36 | Mouse anti-CD9 | Rabbit anti-GFP | 109 | 3 |
| ITGA5-CD36 | Mouse anti GFP | Rabbit anti-ITGA5 | >100 | 4 |
| PTP1B-CD36 | Mouse anti-PTP1B | Rabbit anti-GFP | 90 | 3 |
| CD146-CD36 | Mouse anti-CD146 | Rabbit anti-GFP | 93 | 3 |
| ITGA2-CD36 | Mouse anti-GFP | Rabbit anti-ITGA2 | >100 | 4 |
| ITGB1-CD36 | Mouse anti-Active<br>ITGB1 (12G10) | Rabbit anti-GFP | 69 | 3 |
| Myoferlin-CD36 | Mouse anti-GFP | Rabbit anti-Myoferlin | 73 | 3 |
| CD151-CD36 | Mouse anti-CD151 | Rabbit anti-GFP | 79 | 3 |
| Stomatin-CD36 | Mouse anti-GFP | Rabbit anti-Stomatin | 88 | 3 |
| CD81-CD36 | Mouse anti-CD81 | Rabbit anti-GFP | 72 | 3 |
| ITGA3-CD36 | Mouse anti GFP | Rabbit anti-ITGA3 | >100 | 4 |
| EPCR-CD36 | Mouse anti-EPCR | Rabbit anti-GFP | 55 | 3 |
| Mitofilin-CD36<br>(negative control) | Mouse<br>anti-Mitofilin | Rabbit anti-GFP | 57 | 3 |

### Conditional Colocalization Labeling

TIME HT-CD36 cells were grown to ∼60% confluency on glass coverslips (Electron Microscopy Sciences, Hatfield, PA, USA, ref: 72290-08) and incubated with 0.5 µg/mL of doxycycline for 18 hours prior to immunostaining. JFX549-HaloTag ligand (generous gift from Dr. Lavis, Janelia Research Campus, VA, USA) was diluted to 0.25 nM in MBCD-131 media (Gibco, ThermoFisher, Waltman, MA, USA) and added to cells for 15 minutes at 37°C. For CD36 stimulation experiments, the labelling solution was supplemented with 10 µg/mL of mouse anti-CD36 IgM (clone SMΦ, Santa Cruz Biotechnology, Dallas, TX, USA). Cells were washed and fixed using 4% PFA (Electron Microscopy Sciences, Hatfield, PA, USA) for 15 minutes at room temperature. Following three PBS washes, cells were permeabilized with ice-cold PBS + 0.1% Triton X-100 (Avantor, Radnor, PN, USA) for 10 minutes. Coverslips were subsequently blocked using 3% BSA (Equitech-BioInc, Kerrville, TX, USA) dissolved in PBS for 1 hour at room temperature. Primary antibodies were diluted in PBS with 3% BSA and incubated for 1 hour at room temperature. Antibodies and their concentrations are indicated on Table 5. Cells were washed three times with PBS and incubated with secondary antibodies diluted in PBS with 3% BSA. Coverslips were washed three times in PBS, post-fixed with 4% PFA (Electron Microscopy Sciences, Hatfield, PA, USA) for 10 minutes at room temperature and then washed three times with PBS. Coverslips were imaged via Total Internal Reflection Fluorescence microscopy (TIRF-M).

**Table 5:** List of antibodies and conditions used for conditional colocalization labeling.

| Primary Antibodies | Specificity | Species | Concentra-<br>tion | Provider | Catalog No. |
| --- | --- | --- | --- | --- | --- |
| Anti-CD36<br>(Clone131.2) | CD36 | Mouse | 1:1000 | Gift from Dr.<br>Tandon | N/A |
| Anti-ITGB1 [12G10]<br>AF488 | Active<br>ITGB1 | Mouse | 1:20,000 | Abcam | ab202641 |
| Anti-ITGB1 [12G10]<br>AF647 | Active<br>ITGB1 | Mouse | 1:20,000 | Abcam | ab202643 |
| Anti-CD9<br>[EPR23105-121] | CD9 | Rabbit | 1:10,000 | Abcam | Ab267502 |
| Anti-CD9 AF647<br>[EPR23105-121] | CD9 | Rabbit | 1:10,000 | Abcam | ab267503 |
| Anti-Phospho-SRC<br>(Tyr419) AF488 | pSFK | Rabbit | 1:1000 | Thermo<br>Fisher<br>Scientific | 44-660A1 |
| Anti-TfR (MEM-75)<br>AF647 | TfR | Mouse | 1:10000 | Thermo<br>Fisher<br>Scientific | MA5-18151 |
| Secondary<br>Antibodies | Specificity | Species | Concentra-<br>tion | Provider | Catalog No. |
| Donkey Anti-Rabbit<br>AF 488 | Rabbit IgG | Donkey | 1:500 | Jackson<br>Immuno<br>Research | 711-545-152 |

### CD36 stimulation and pSFK immunostaining

The levels of Fyn activation were determined using an antibody directed against the phosphorylated tyrosine 420 (P-Y420) residue of Src family kinases. First, cells were serum starved for 3 hours in MBCD-131 media (Gibco, ThermoFisher, Waltman, MA, USA). Following serum starvation, cells were stimulated with MBCD-131 containing 10 µg/mL mouse anti-IgM (clone SMΦ, Santa Cruz Biotechnology, Dallas, TX, USA) for 15 minutes. Cells were then processed for immunostaining or conditional colocalization labeling. For immunostaining, cells were permeabilized and block simultaneously (PBS with 0.1% TX-100 and 3% BSA) for 30 min at room temperature, before being incubated with rabbit anti-P-Y420-Src (pSFK) antibody coupled to AF488 (ThermoFisher, 44-660A1) and phalloidin-AF568 (ThermoFisher, ref: A12380). Coverslips were imaged by TIRFm.

### Fluorescence microscopy imaging

Regular widefield imaging was performed on an Olympus IX81 stand (Evident Canada Inc., Quebec, QC, Canada) equipped with Diskovery module (Quorum Technologies, Guelph, ON, Canada). Fluorescent excitations were from a X-Cite 120 light source (Excelitas Technologies Corp, Pittsburgh, PA USA) and the appropriate dichroic filters (DAPI, GFP or Rhodamine cubes). Image acquisitions were performed with an oil immersion 60×1.49 numerical aperture objective on an ImageEM Hamamatsu EM-CDD camera (ImageEM91013, Hamamatsu Photonics, Japan) using Volocity software (Quorum Technologies, Guelph, ON, Canada).

Confocal imaging was carried out on a Diskovery system (Quorum Technologies, Guelph, ON, Canada) set-up on an Olympus IX81 stand (Evident Canada Inc., Quebec, QC, Canada) in spinning disk mode, which consisted of inserting a rotating 50 µm pinhole disk into the light path. Fluorescent dyes were excited using a laser merge module from Spectral Applied Physics (Richmond Hill, On, Canada) equipped with 491 nm, 561 and 643 nm solid-state lasers and a multi-mode laser randomizer. Image acquisitions were performed using the appropriate emission filters (Chroma, Bellows Falls, VT, USA) mounted on a filter-wheel (Sutter Instrument, Novato, CA, USA) through an oil immersion 60x 1.49 numerical aperture objective with an ImageEM Hamamatsu EM-CDD camera (ImageEM91013, Hamamatsu Photonics, Japan) using Volocity software (Quorum Technologies, Guelph, ON, Canada).

Total Internal Reflection Fluorescence microscopy (TIRFm) images were captured on an Olympus IX-81 motorized inverted TIRF base (Evident Canada Inc., Quebec, QC, Canada) installed by Quorum Technologies (Guelph, ON, Canada). Fluorescent dyes were excited using laser merge module from Spectral Applied Physics (Richmond Hill, On, Canada) equipped with 491 nm, 561 and 643 nm solid-state lasers and a single-mode laser attached to the Evident TIRF illuminator, with incident light angle adjustable manually. Images were recorded using the appropriate emission filters (Chroma, Bellows Falls, VT, USA) mounted on a filter-wheel (Sutter Instrument, Novato, CA, USA) with a Hamamatsu EM-CDD camera (ImageEM91013, Hamamatsu Photonics, Japan) using Volocity software (Quorum Technologies, Guelph, ON, Canada) through an 100×1.49 NA objective.

### Quantification of PLA Dot Density per Cell

PLA images taken in confocal mode were loaded into CellProfiler (Stirling et al., 2021). These three colour TIFF images were composed of the DAPI (blue channel), concanavalin-A AF488 (green channel), and PLA signal (far-red channel). The blue, green and far-red channels were split into grayscale images using the ‘colortogreymodule’. Segmentation of each cell was performed using concanavalinA-AF488 cell mask and the ‘IdentifyObjectManually’ module. The cell masks obtained from the ‘IdentifySecondaryObject’ were converted to binary images. The integrated intensities for the binary images were determined using the ‘MeasureObjectIntensity’ module. Pixels containing the cell masks were given a value of 1 in the binary image, and therefore the integrated intensities allowed us to determine the area, in pixels, of each cell.

The module ‘IdentifyPrimaryObject’ was utilized to segment and count the number of PLA dots in each image. Minimum Cross-Entropy thresholding was used for segmenting PLA dots. Minimum Cross-entropy algorithm assigns a thresholding value for each pixel, which minimizes the probability of misclassifying the background or foreground. The number of PLA per cell was determined using the cell masks obtained from the ‘IdentifySecondaryObject’ module.

Segmented images were saved for review using the ‘Save Images’ module. Finally, the number of PLA dots per cell and the area of each cell were exported using the ‘ExportToSpreadsheet’ module. To normalize for cell size, the density of PLA dots per cell was calculated by dividing the number of PLA dots found in a cell to its area in pixel. For statistical comparisons of PLA experiments, a Krushkal-Wallis non-parametric ANOVA test was performed followed by a Dunn’s post-hoc test (Kruskal and Wallis, 1952; Dunn, 1964). P-values were adjusted using Bonferroni correction (Etymologia: Bonferroni Correction, 2015). An α-value of 0.05 or less was chosen for significance.

### Conditional Colocalization Analysis

Images obtained by TIRFm for cells prepared for conditional colocalization staining were analyzed using the conditional colocalization package developed in MATLAB (MathWorks, Natick, MA, USA) by Vega-Lugo et al. (Vega-Lugo et al., 2022) following the instructions provided (https://github.com/kjaqaman/conditionalColoc). Three color TIFF images were saved as a movie list. Channels were assigned to three groups: ‘target,’ ‘reference,’ or ‘condition’. The ‘point-source detection’ algorithm in u-track (https://github.com/DanuserLab/u-track) was used to determine the subpixel x,y coordinates of each punctate objects (see Table 6 for a summary of the parameters used for detection). Objects were defined to colocalize if the distance between the centers of two punctate objects was less than 2 pixels (∼180 nm). The region of interest defining the cell area on each image was drawn manually. Wilcoxon rank sum test was used to determine the statistical significance (Pratt, 1959). Thresholds for three molecular colocalization significance were calculated using Dunn-Sidak correction (Dunn, 1964; Sidak, 1967). To learn more about the conditional colocalization analysis, refer to Vega-Lugo et al. (Vega-Lugo et al., 2022). For statistical comparison between conditional colocalization values between unstimulated and stimulated treatments a non-parametric T-test was performed (Welch, 1947). An α-value of 0.05 was chosen for significance for all conditional colocalization comparisons.

**Table 6:**
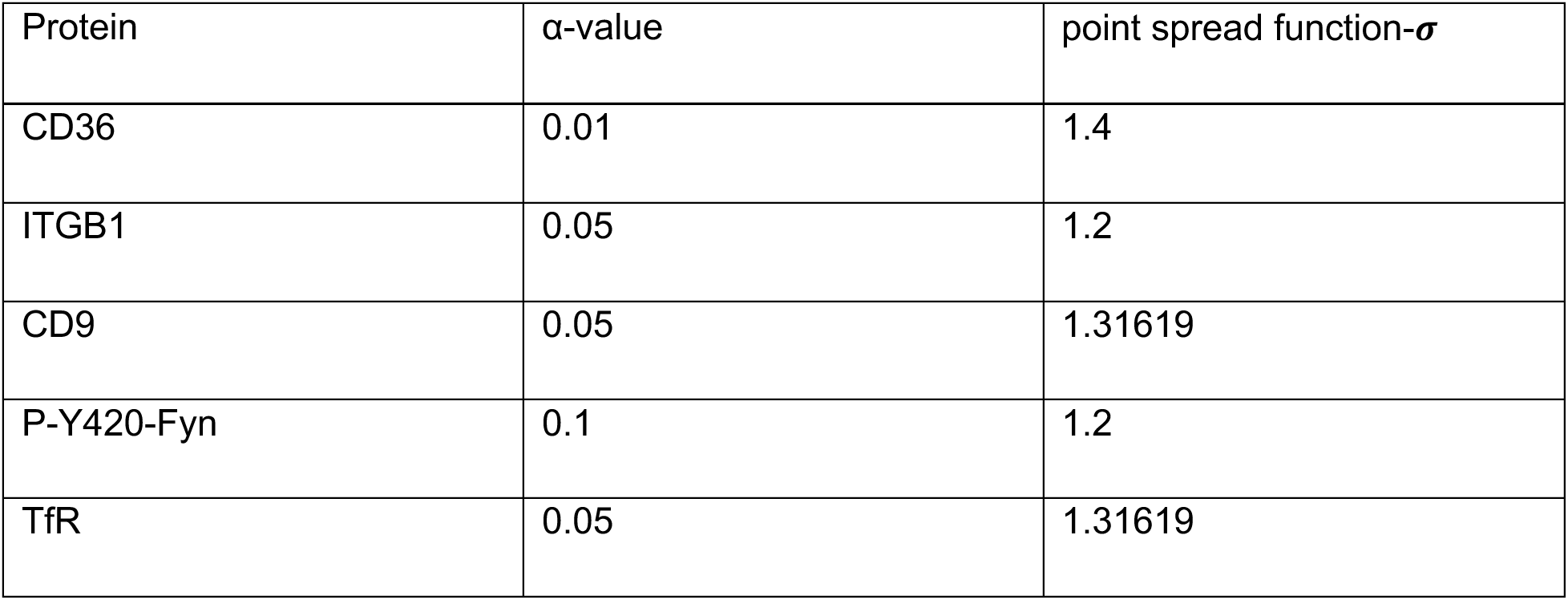
Detection parameters for conditional colocalization analysis. The α-value and point spread function-# used for point source detection of CD36, ITGB1, CD9, P-Y420-Fyn, and TfR.

|  |  |  |
| --- | --- | --- |
| Protein | α-value | point spread function-σ |
| CD36 | 0.01 | 1.4 |
| ITGB1 | 0.05 | 1.2 |
| CD9 | 0.05 | 1.31619 |
| P-Y420-Fyn | 0.1 | 1.2 |
| TfR | 0.05 | 1.31619 |

### Quantification of pSFK intensity

TIRFm images of F-actin and pSFK were acquired with similar parameters across all experiments. Images were loaded in customized quantification pipeline developed in CellProfiler (Stirling et al., 2021). Two color TIFF 16-bit images were composed of the phalloidin-AF568 (red channel) and pSFK (green channel). The channels were split into grayscale images using the ‘colortogreymodule’. Segmentation of each cell was performed using phalloidin AF568 channel and the ‘IdentifyObjectManually’ module. The area of the cell masks obtained from ‘IdentifyObjectManually’ were measured using the “MeasureImageAreaOccupied”. To determine the total intensity of pSFK, we inputted the pSFK (green channel) into the “MeasureImageIntensity” module. Finally, the total intensity of activated Fyn and the area of each cell were exported using the ‘ExportToSpreadsheet’ module. The mean intensity of pSFK was calculated by dividing the total pSFK intensity by the cell area in pixel. For statistical comparisons of CD36 stimulation experiments, a non-parametric T-test was performed (Welch, 1947). An α-value of 0.05 or less was chosen for significance.

### Graphical Representation and Statistical Analysis

Data from PLA, conditional colocalization, and CD36-IgM stimulation experiments were displayed using boxplots. The median of the data set is represented by the central line. Above and below the median line are notches indicating the 95% confidence interval of the median line. The edges of the boxplot represent the 25^th^ and 75^th^ percentiles and the whiskers display the maximum and minimum data points that were not outliers. Data points outside the whiskers of the boxplot are outliers, and usually shown in red. For PLA, CD36 stimulation, and conditional colocalization experiments statistical and graphical analysis were performed using SciPy (Virtanen et al., 2020) or MATLAB (MathWorks, Natick, MA, USA). Graphing and statistics of conditional colocalization data were performed using MATLAB (MathWorks, Natick, MA, USA). Figures were generated in Adobe Illustrator (Adobe, San Jose, CA, USA).

## Data availability

All MS raw data files were uploaded to the publicly accessible mass spectrometry repository MassIVE (https://massive.ucsd.edu, MSV000094847).

## Acknowledgements

We would like to thank Touret, Julien and Fahlman lab members for helpful discussions and suggestions. We thank Jack Moore and the Alberta Proteomics and Mass Spectrometry facility for their help. We also thank Kiara Smith, Dr. Strickfaden and the Cell Imaging Centre for their assistance with microscopy.

This work was supported by an infrastructure CFI-JELF award (O.J. 37833 and 39051), and operating grants from the Natural Sciences and Engineering Research Council of Canada (O.J. RGPIN-2018-05881 and N.T. RGPIN-2018-05783), Canadian Institutes for Health Research (N. T. PS 165816), National Science Foundation (K.J. MCB-2114417), the National Institutes of Health/National Institute of General Medical Sciences (KJ R35 GM119619), and the UTSW Endowed Scholars Program (K.J.). EGC was supported by the Canadian Glycomics Network Strategic Science Fund (RES0063541).

## Abbreviations

BAR: Biotinylation by Antibody Recognition
EC: endothelial cells
EPCR: Endothelial Protein Receptor C
FT: Flow-Through
ITGB1: Integrin Beta-1
HDMEC: Human Dermal Microvascular Endothelial Cells
HT: HaloTag
KD: knockdown
mEm: mEmerald fluorescent protein
MYOF: Myoferlin
PL: Proximity Labeling
PLA: Proximity Ligation Assay
STOM: Stomatin
TIME: Human Telomerase-immortalized Microvascular Endothelial Cells
WCL: Whole Cell Lysate.

