## Supplementary figures for "Investigation of CD36 interactome provides insights into multimolecular complexes necessary for anti-angiogenic signalling"

#### Supplementary Figure 1

**A**

##### CD36 BAR experiments

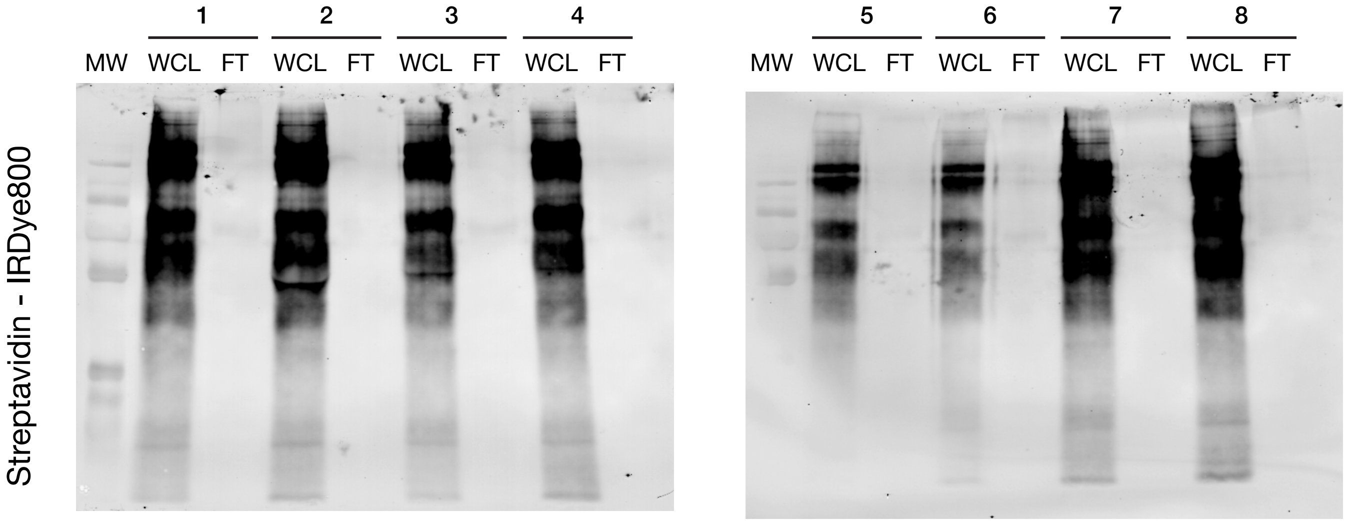


**B**

##### BAR control experiments

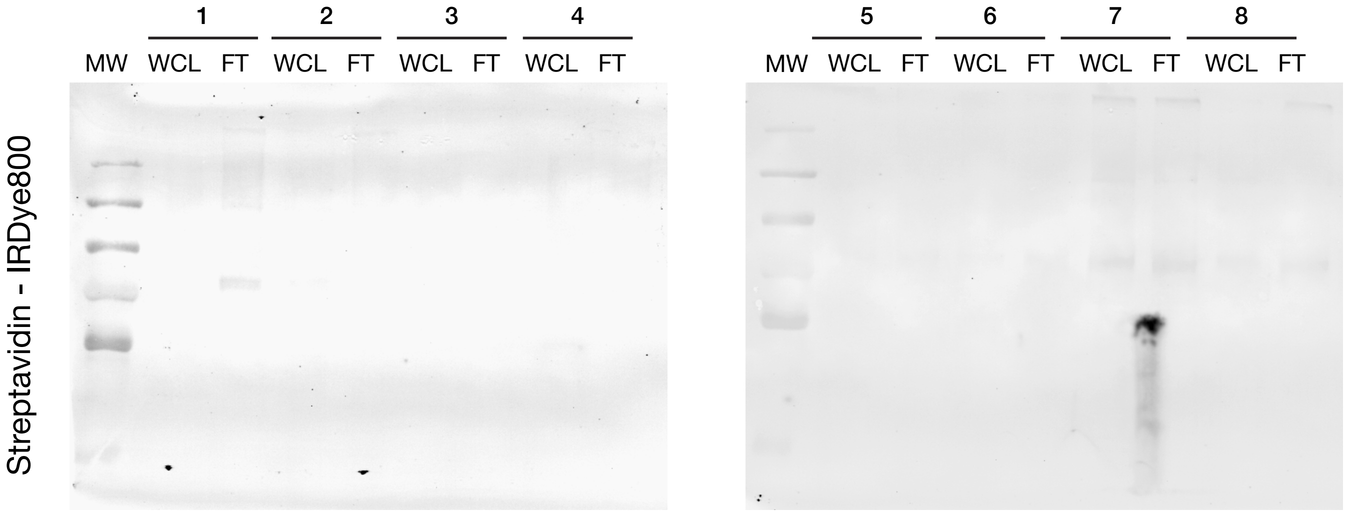

**Supplementary Figure 1: Representative streptavidin blots of Whole Cell Lysates and Flow Through after biotin-capture for BAR experiments.**

TIME mEm-CD36 cells were grown in 6 cm dishes and (**A**) induced (CD36 BAR) with doxycycline for 18 hours or (**B**) not induced (BAR control) before fixation and BAR experiments (Fig. 1A and materials and methods). About 10% vol. of Whole Cell Lysates (WCL) was kept for evaluation of biotinylation reactions while the remaining 90% was processed for capture via streptavidin beads, trypsinization and peptide identification via LC-MS/MS. Flow through (FT) correspond to supernatants recovered after capture of biotinylated proteins. Both WCL and FT were run on SDS-PAGE, transfer to nitrocellulose and probed with streptavidin-IRDye 800RD.

### Supplementary Figure 2

**A**

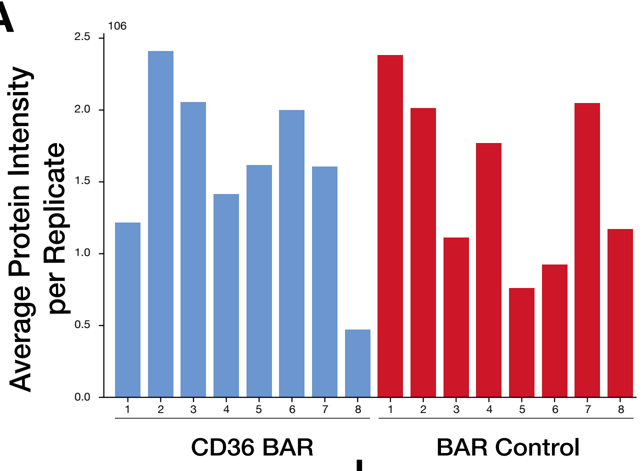


367 common proteins found across all 16 samples

| Samples | Average common protein intensity ($10^6$) | Normalization Factor |
|---|---|---|
| CD36 BAR - 1 | 4.16 | 2.69 |
| CD36 BAR - 2 | 8.72 | 5.64 |
| CD36 BAR - 3 | 6.57 | 4.25 |
| CD36 BAR - 4 | 4.51 | 2.92 |
| CD36 BAR - 5 | 5.74 | 3.71 |
| CD36 BAR - 6 | 7.30 | 4.72 |
| CD36 BAR - 7 | 5.83 | 3.77 |
| CD36 BAR - 8 | 1.55 | 1.00 |
| BAR Control - 1 | 8.51 | 5.50 |
| BAR Control - 2 | 7.16 | 4.62 |
| BAR Control - 3 | 3.95 | 2.55 |
| BAR Control - 4 | 6.32 | 4.09 |
| BAR Control - 5 | 2.69 | 1.74 |
| BAR Control - 6 | 3.31 | 2.14 |
| BAR Control - 7 | 4.65 | 3.00 |
| BAR Control - 8 | 4.27 | 2.76 |

**B**

CD36 BAR Mass Spectrometry datasets

Determination of Normalization factors

Find zero values

| Protein Names | BAR - 1 | BAR - 2 | ... | Control - 1 | Control - 2 |
|---|---|---|---|---|---|
| Galactokinase | 0.00 | 0.00 | ... | $2.89*10^4$ | $1.88*10^4$ |
| 40s ribosomal protein S3a | $1.70*10^5$ | $1.29*10^6$ | ... | $1.30*10^8$ | $2.03*10^8$ |
| Calnexin | $7.33*10^4$ | $2.91*10^5$ | ... | 0.00 | 0.00 |
| Acyl-protein thioesterase 2 | $5.45*10^5$ | $6.81*10^5$ | ... | 0.00 | $5.37*10^3$ |
| ... | ... | ... | ... | ... | ... |
| Minimum protein Intensity for Sample | $2.70*10^3$ | $2.08*10^3$ | ... | $1.24*10^3$ | $2.57*10^3$ |

Replace 0 with lowest intentity value of the condition

| Protein Names | BAR - 1 | BAR - 2 | ... | Control - 1 | Control - 2 |
|---|---|---|---|---|---|
| Galactokinase | $2.70*10^3$ | $2.08*10^3$ | ... | $2.89*10^4$ | $1.88*10^4$ |
| 40s ribosomal protein S3a | $1.70*10^5$ | $1.29*10^6$ | ... | $1.30*10^8$ | $2.03*10^8$ |
| Calnexin | $7.33*10^4$ | $2.91*10^5$ | ... | $1.24*10^3$ | $2.57*10^3$ |
| Acyl-protein thioesterase 2 | $5.45*10^5$ | $6.81*10^5$ | ... | $1.24*10^3$ | $5.37*10^3$ |
| ... | ... | ... | ... | ... | ... |
| Minimum protein Intensity for Sample | $2.70*10^3$ | $2.08*10^3$ | ... | $1.24*10^3$ | $2.57*10^3$ |

Applying Normalization factor for each condition

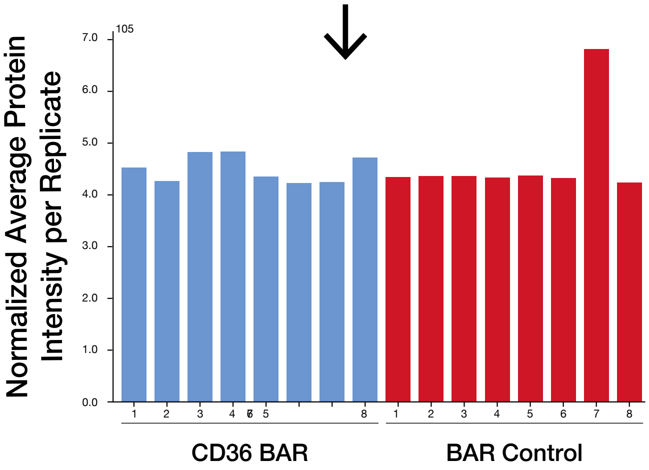


**C**

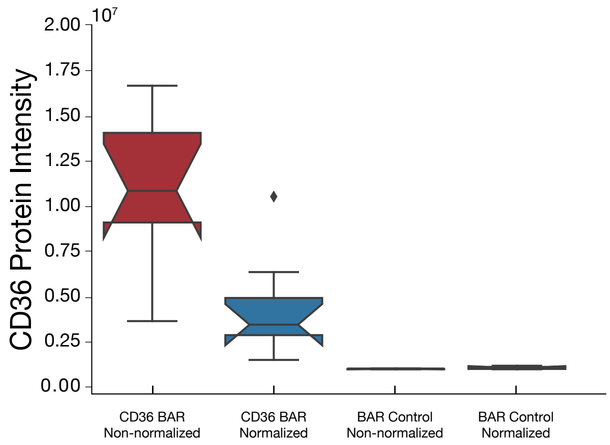

**Supplementary Figure 2: Normalization workflow of CD36 BAR mass spectrometry data protein intensity.**

(**A**) Determination of the normalization factor for each CD36 BAR and BAR control samples. 367 common proteins were identified between all conditions, and the average protein intensity for these common proteins was calculated. Normalization factors were calculated by dividing the minimum average protein intensity (indicated in green) by the average protein intensity for the BAR samples. (**B**) Normalization of CD36 BAR mass spectrometry data. Before normalization, zero-filling was performed by replacing blank values with the minimum protein intensity identified within each sample (local minima). Normalization of protein intensities was obtained by dividing all protein intensities within a sample by their respective normalization factor. (**C**) CD36 normalized and non-normalized protein intensity within CD36 BAR and BAR control datasets.

#### Supplementary Figure 3

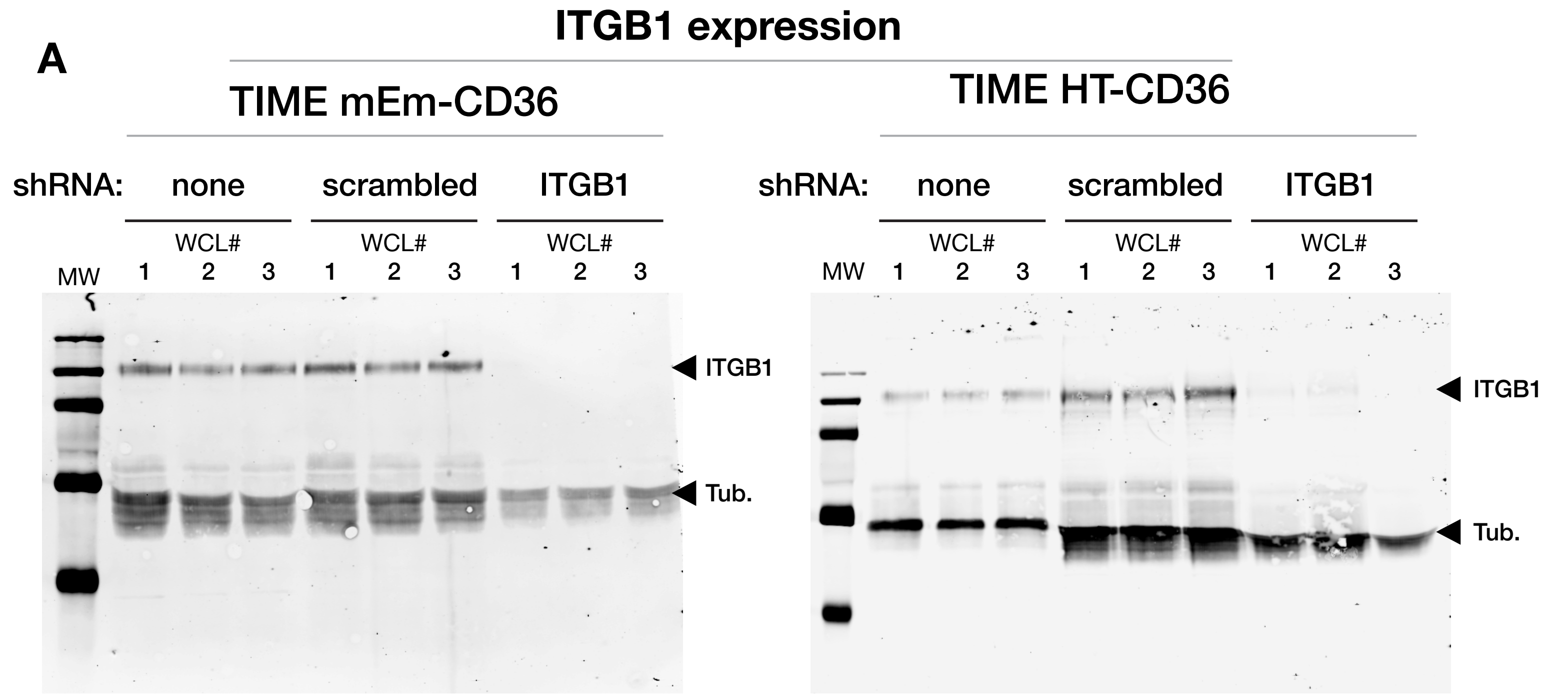


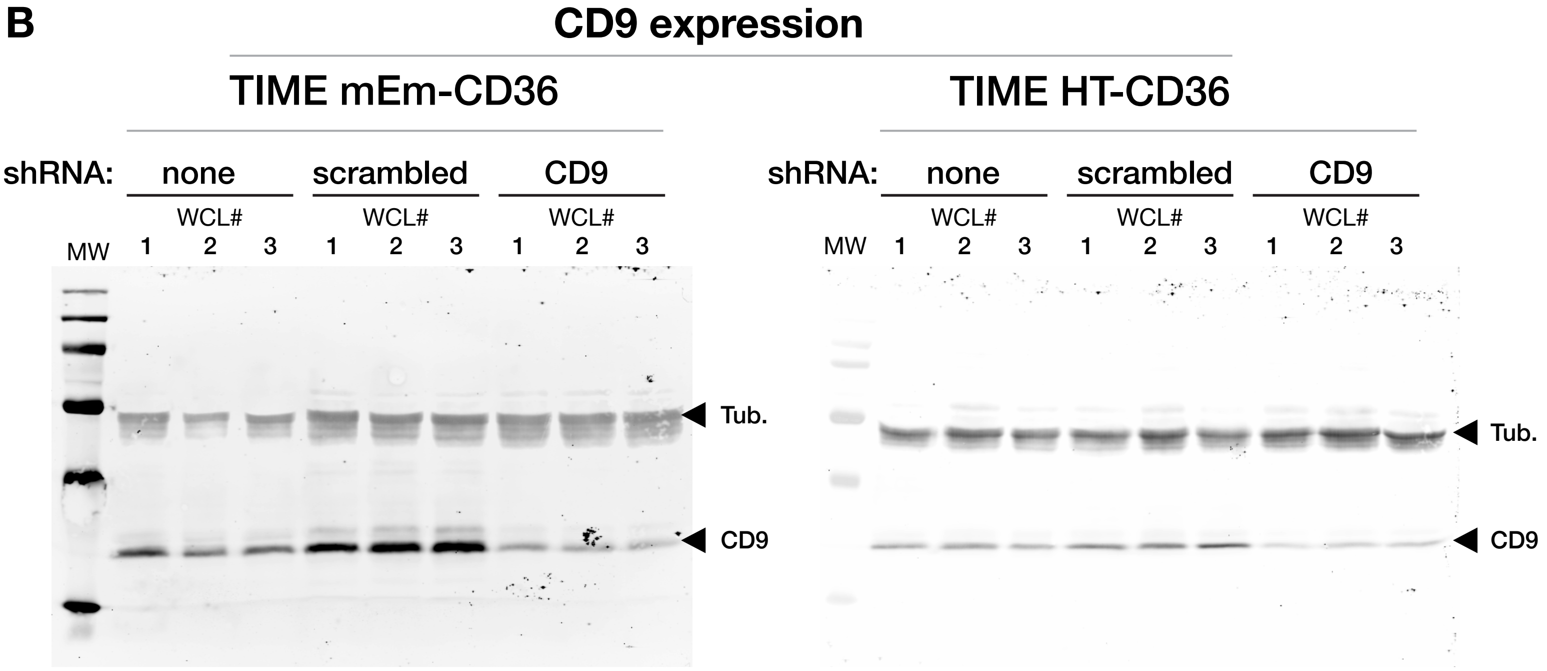

**Supplementary Figure 3: Validation of TIME mEmerald-CD36 and HaloTag-CD36 ITGb1 and CD9 shRNA KD via Immunoblotting.**

TIME mEm-CD36 and TIME HaloTag-CD36 cells were transduced with lentiviruses containing sequences for the constitutive expression of scrambled, ITGB1, or CD9 shRNA. Following hygromycin selection, surviving cells were collected and their lysate (WCL from 3 replicates) was analyzed via immunoblotting to confirm candidate protein silencing. (**A**) Immunoblots evaluating the expression of ITGB1 in TIME mEm-CD36 and TIME HT-CD36 cell lines, and cell lines expressing ITGB1 or scrambled shRNA. Arrows on the side indicate suggested molecular weight for ITGB1 and tubulin. (**B**) Immunoblots evaluating the expression of CD9 in TIME mEmerald-CD36 and TIME HT-CD36 cell lines, and cell lines expressing CD9 or scrambled shRNA. Arrows on the side indicate suggested molecular weight for CD9 and tubulin. For Fig. 4A, and B, the expression levels of ITGB1 and CD9 were quantified with Empira Studio Software (Li-Cor Bioscience) and normalized to tubulin loading control.

**Supplementary Table 1: List of candidate proteins enriched and significant in CD36 BAR.**

| Protein Name | Ratio | p-value |
|---|---|---|
| Neutral alpha-glucosidase AB | 179.356 | <0.001 |
| 40S ribosomal protein S17 | 149.85 | 0.007 |
| Disintegrin and metalloproteinase domain-containing protein 10 | 148.088 | 0.001 |
| Platelet glycoprotein 4 | 140.081 | 0.014 |
| Synaptosomal-associated protein 23 | 131.331 | 0.019 |
| Fibulin-1 | 126.475 | 0.005 |
| Poliovirus receptor | 122.905 | 0.037 |
| CD81 antigen | 121.593 | 0.010 |
| Junctional adhesion molecule A | 112.754 | 0.028 |
| 39S ribosomal protein L38, mitochondrial | 105.483 | 0.033 |
| Cytochrome c oxidase subunit 2 | 96.256 | <0.001 |
| Protein disulfide-isomerase A6 | 94.529 | 0.002 |
| Acyl-protein thioesterase 2 | 92.353 | 0.002 |
| Endothelial protein C receptor | 89.145 | 0.010 |
| Carbonyl reductase [NADPH] 1 | 84.315 | 0.015 |
| Survival motor neuron protein | 75.082 | 0.011 |
| UTP--glucose-1-phosphate uridylyltransferase | 74.208 | 0.016 |
| GTPase KRas | 57.731 | 0.011 |
| Collagen alpha-1(XVIII) chain | 53.486 | 0.001 |
| Heterogeneous nuclear ribonucleoprotein H2 | 52.501 | 0.020 |
| C-type lectin domain family 14 member A | 51.218 | 0.026 |

| | | |
|---|---|---|
| Cytoplasmic aconitate hydratase | 47.163 | 0.039 |
| Endothelial cell-selective adhesion molecule | 42.82 | 0.004 |
| Elongation factor 1-alpha 1 | 36.256 | 0.005 |
| Neuropilin-1 | 35.567 | 0.008 |
| Calcium homeostasis endoplasmic reticulum protein | 34.354 | 0.002 |
| Nectin-2 | 34.069 | 0.030 |
| Peptidyl-prolyl cis-trans isomerase FKBP8 | 33.818 | 0.019 |
| ADAMTS-like protein 1 | 31.546 | 0.008 |
| Rab GTPase-activating protein 1 | 30.341 | 0.005 |
| Serine/threonine-protein kinase N2 | 29.76 | 0.005 |
| cAMP-dependent protein kinase type I-alpha regulatory subunit | 29.452 | 0.018 |
| Collagen alpha-1(VIII) chain | 29.311 | 0.024 |
| Coagulation factor V | 28.004 | 0.007 |
| E3 ubiquitin-protein ligase HUWE1 | 27.305 | 0.004 |
| Intercellular adhesion molecule 2 | 26.437 | 0.020 |
| Calreticulin | 26.087 | 0.016 |
| Protein Niban 2 | 22.427 | 0.012 |
| Integrin alpha-5 | 22.356 | 0.001 |
| CD59 glycoprotein | 20.217 | 0.001 |
| Multidrug resistance-associated protein 1 | 19.265 | 0.007 |
| EF-hand domain-containing protein D2 | 18.24 | 0.003 |
| Peroxiredoxin-2 | 18.16 | 0.040 |
| Lon protease homolog, mitochondrial | 17.529 | 0.018 |

| | | |
|---|---|---|
| Integrin alpha-3 | 16.591 | <0.001 |
| Integrin beta-1 | 15.138 | 0.004 |
| Serum albumin | 15.128 | 0.015 |
| 28S ribosomal protein S25, mitochondrial | 14.916 | 0.026 |
| Endoglin | 14.862 | 0.002 |
| Tyrosine-protein kinase Fyn | 14.65 | 0.009 |
| Erythrocyte band 7 integral membrane protein | 14.616 | 0.022 |
| Plexin-D1 | 14.541 | 0.002 |
| Integrin alpha-6 | 14.346 | 0.015 |
| Splicing factor 3A subunit 3 | 14.236 | 0.009 |
| Aminopeptidase N | 13.488 | 0.004 |
| Double-stranded RNA-binding protein Staufen homolog 1 | 13.216 | 0.021 |
| Disintegrin and metalloproteinase domain-containing protein 9 | 13.198 | 0.034 |
| Y-box-binding protein 1 | 12.518 | 0.027 |
| Laminin subunit gamma-1 | 12.015 | 0.038 |
| AP-3 complex subunit delta-1 | 11.786 | 0.002 |
| Sushi repeat-containing protein SRPX | 11.045 | 0.039 |
| Laminin subunit alpha-4 | 10.593 | 0.033 |
| CD9 antigen | 10.585 | 0.014 |
| Transforming growth factor-beta-induced protein ig-h3 | 10.523 | 0.012 |
| Thrombospondin type-1 domain-containing protein 4 | 10.326 | 0.007 |
| Replication factor C subunit 2 | 10.225 | 0.011 |
| Flotillin-2 | 9.653 | 0.005 |

| | | |
|---|---|---|
| HLA class I histocompatibility antigen, C alpha chain | 9.294 | <0.001 |
| Annexin A6 | 8.86 | 0.027 |
| Protein transport protein Sec24A | 8.437 | 0.004 |
| Hemoglobin subunit alpha | 8.32 | 0.017 |
| NEDD8-conjugating enzyme Ubc12 | 8.229 | 0.040 |
| Microtubule-actin cross-linking factor 1, isoforms 1/2/3/5 | 8.19 | 0.025 |
| Fibrillin-1 | 8.087 | 0.007 |
| Guanine nucleotide-binding protein G(i) subunit alpha-2 | 8.032 | 0.005 |
| EGF-containing fibulin-like extracellular matrix protein 1 | 7.485 | 0.001 |
| HLA class I histocompatibility antigen, B alpha chain | 7.482 | 0.029 |
| Rab GDP dissociation inhibitor beta | 7.43 | 0.003 |
| Latent-transforming growth factor beta-binding protein 1 | 7.238 | 0.034 |
| Mitochondrial-processing peptidase subunit alpha | 7.045 | 0.032 |
| Isoform 2 of CD166 antigen | 6.999 | 0.019 |
| Ran GTPase-activating protein 1 | 6.981 | 0.013 |
| TAR DNA-binding protein 43 | 6.791 | 0.046 |
| CD109 antigen | 6.733 | 0.005 |
| Thioredoxin-dependent peroxide reductase, mitochondrial | 6.666 | 0.015 |
| 26S proteasome non-ATPase regulatory subunit 3 | 6.655 | 0.002 |
| Desmoglein-1 | 6.602 | 0.001 |
| Platelet endothelial cell adhesion molecule | 6.461 | <0.001 |
| Ephrin type-A receptor 2 | 6.411 | 0.005 |
| Developmentally-regulated GTP-binding protein 1 | 6.326 | 0.041 |

| | | |
|---|---|---|
| Integrin alpha-2 | 6.254 | 0.002 |
| Ras-related protein Rab-7a | 6.234 | 0.046 |
| Catenin alpha-1 | 6.19 | 0.016 |
| Breakpoint cluster region protein | 6.167 | 0.012 |
| Guanine nucleotide-binding protein subunit beta-4 | 6.084 | 0.016 |
| Activating signal cointegrator 1 complex subunit 2 | 5.943 | 0.048 |
| Isoform 4 of Leucine-rich repeat flightless-interacting protein 1 | 5.911 | 0.019 |
| Calponin-2 | 5.878 | 0.030 |
| DNA-directed RNA polymerase II subunit RPB3 | 5.873 | 0.043 |
| Plasminogen activator inhibitor 1 | 5.725 | 0.018 |
| CD44 antigen | 5.715 | 0.009 |
| Src substrate cortactin | 5.485 | 0.034 |
| DBIRD complex subunit ZNF326 | 5.261 | 0.011 |
| Signal recognition particle subunit SRP68 | 5.155 | 0.018 |
| Eukaryotic translation initiation factor 3 subunit A | 4.856 | 0.002 |
| Antithrombin-III | 4.815 | 0.029 |
| Importin subunit alpha-7 | 4.711 | 0.041 |
| Tyrosine-protein phosphatase non-receptor type 1 | 4.695 | 0.019 |
| Dolichyl-diphosphooligosaccharide--protein glycosyltransferase subunit 1 | 4.634 | 0.005 |
| U3 small nucleolar RNA-associated protein 14 homolog A | 4.63 | 0.013 |
| Junction plakoglobin | 4.609 | 0.009 |
| WASH complex subunit 4 | 4.512 | 0.011 |

| | | |
|---|---|---|
| Isocitrate dehydrogenase [NADP], mitochondrial | 4.359 | 0.008 |
| E3 ubiquitin-protein ligase ARIH1 | 4.354 | 0.039 |
| Multimerin-1 | 4.338 | 0.038 |
| Peptidyl-prolyl cis-trans isomerase FKBP1A | 4.273 | 0.012 |
| Cadherin-5 | 4.262 | 0.005 |
| Basigin | 4.192 | 0.048 |
| Desmoplakin | 4.101 | 0.015 |
| Voltage-dependent anion-selective channel protein 1 | 3.954 | 0.036 |
| 60S ribosomal protein L23 | 3.842 | 0.015 |
| Pyruvate kinase PKM | 3.76 | 0.003 |
| Malate dehydrogenase, mitochondrial | 3.701 | 0.008 |
| Squamous cell carcinoma antigen recognized by T-cells 3 | 3.573 | 0.048 |
| Glutamate dehydrogenase 1, mitochondrial | 3.495 | 0.003 |
| Histone-arginine methyltransferase CARM1 | 3.317 | 0.004 |
| Nuclear pore complex protein Nup133 | 3.305 | 0.043 |
| Multimerin-2 | 3.304 | 0.035 |
| Dihydrolipoyl dehydrogenase, mitochondrial | 3.286 | <0.001 |
| Citrate synthase, mitochondrial | 3.26 | 0.013 |
| Leucine-rich PPR motif-containing protein, mitochondrial | 3.256 | 0.026 |
| A-kinase anchor protein 12 | 3.194 | 0.001 |
| 60S ribosomal protein L18a | 3.11 | 0.022 |
| Myoferlin | 3.055 | 0.023 |
| AP2-associated protein kinase 1 | 3.036 | 0.045 |

| | | |
|---|---|---|
| Hexokinase-1 | 2.928 | 0.006 |
| Prothrombin | 2.921 | 0.024 |
| Trifunctional enzyme subunit alpha, mitochondrial | 2.871 | 0.006 |
| 5'-nucleotidase | 2.859 | 0.044 |
| Peroxidasin homolog | 2.761 | 0.033 |
| 60S ribosomal protein L37a | 2.704 | 0.030 |
| Catenin delta-1 | 2.673 | 0.001 |
| ATP-binding cassette sub-family F member 1 | 2.519 | 0.031 |
| Ras GTPase-activating-like protein IQGAP1 | 2.461 | 0.008 |
| Moesin | 2.426 | 0.022 |
| Ankycorbin | 2.329 | 0.018 |
| EH domain-containing protein 2 | 2.3 | 0.040 |
| Annexin A2 | 2.267 | <0.001 |
| 40S ribosomal protein SA | 2.186 | 0.021 |
| Cell surface glycoprotein MUC18 | 2.173 | 0.048 |
| 60S ribosomal protein L19 | 2.084 | 0.020 |
| Ras-interacting protein 1 | 2.021 | 0.013 |
| EH domain-containing protein 4 | 2.001 | 0.045 |
